# Low dimensional latent structure underlying the choices of mice

**DOI:** 10.1101/2025.08.08.669381

**Authors:** Sebastian A. Bruijns, Maria K. Eckstein, Peter Dayan

**Affiliations:** Max Planck Institute for Biological Cybernetics, Tübingen, Germany; University of Tübingen, Germany; Google DeepMind

## Abstract

An impressive wealth of cognitive neuroscience tasks involves combining perceptual information with an estimate of the latent state of the environment to make a decision. Such tasks have driven the development of theoretically-motivated cognitive models which offer compactly parameterised, and thereby insightful, accounts of the internal processes by which this might happen. However, the very large amounts of data that can now be collected present a challenge and an opportunity for this framework. The challenge is that these models can be shown to underfit the data systematically (Nassar & Frank, 2016; Palminteri, Wyart, & Koechlin, 2017; Wilson & Collins, 2019) – particularly in how they characterise the effect of the latent state estimate. But this opens up the opportunity to extend the framework and employ richer, more highly parameterised and flexible models, in a data-driven manner (Dezfouli, Griffiths, Ramos, Dayan, & Balleine, 2019). Unfortunately, it is tremendously difficult to interpret these models, precisely because of their flexibility. Here, we follow a recent approach (Eckstein, Summerfield, Daw, & Miller, 2024), in which components of compact models are progressively replaced with more flexible homologues, but, where possible, the resulting insights are mapped back into theoretically-transparent forms. We prove the effectiveness of our scheme by applying it to one of the largest available decision-making data sets for mice – more than 300,000 choices from 139 subjects studied by the International Brain Lab (The International Brain Laboratory et al., 2021). We found widely generalisable phenomena such as notable effects of continual fluctuations in task engagement and systematic differences between learning and forgetting of chosen and unchosen options in determining the subjective state estimate. Our results show that combining theory-driven and data-driven methods can reveal cognitive processes that would have been difficult to discover using either method on its own.

---

The International Brain Laboratory (IBL; The International Brain Laboratory et al., 2021) administered a paradigmatic decision-making task in which head-fixed mice were shown a sinusoidal grating of a controlled contrast on either the right or left side of a screen, accompanied by a tone that signified the beginning of a trial. Mice then had to center the grating (within 60 s) by turning a steering wheel in the appropriate direction (see **Fig. 1a**). Successful trials led to water reward; unsuccessful trials to a noise burst and a 1 s timeout. We considered expert behaviour on this task, which follows on from a rigorous shaping procedure (that is also amenable to modelling; Bruijns et al. 2023).

**Fig. 1.**
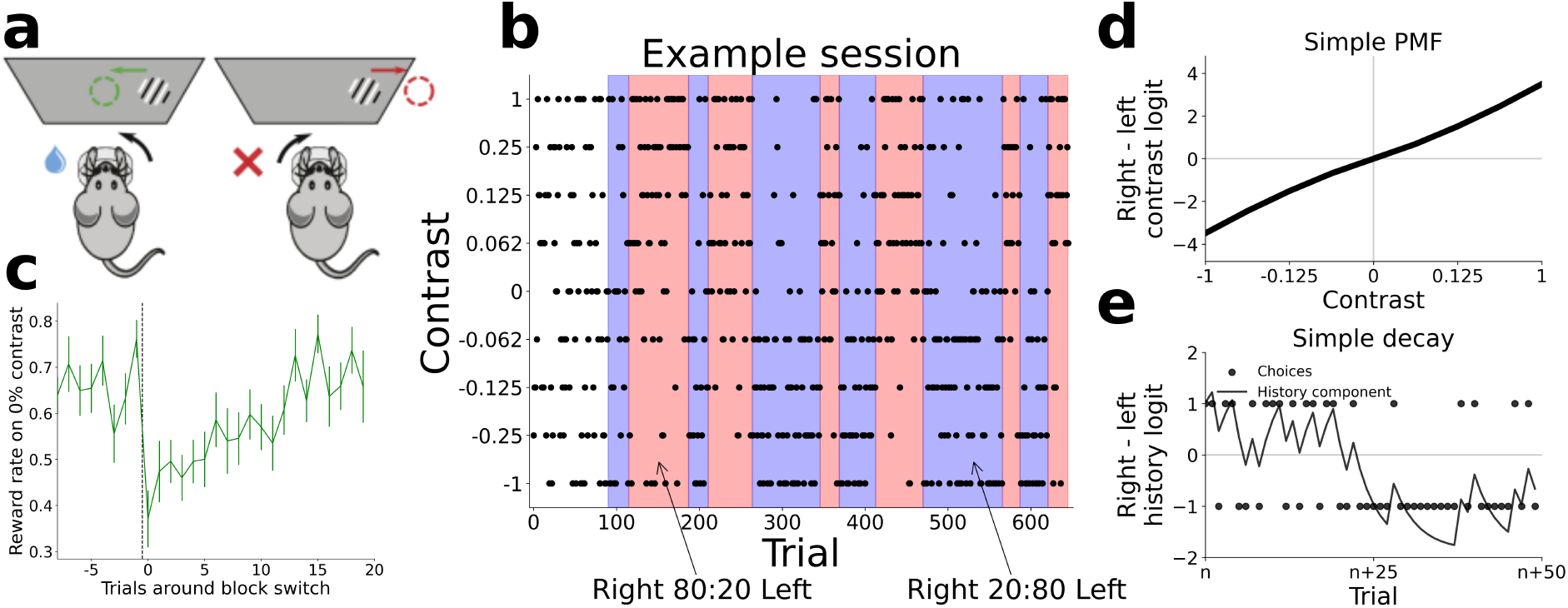
The IBL task and mouse behaviour. **(a)** Left: The mouse performs a rightwards action (turning the wheel counterclockwise), thereby centering the stimulus originally presented on the righthand side of the screen and earning a water reward. Right: A clockwise turn (the leftwards choice) would move the stimulus outside the screen, resulting in a noise burst and a timeout before the onset of the next trial. **(b)** An example session, showing which contrast was shown on which trial, and which block is currently active (background colour: white, neutral first 90 trials; blue, left biased; red, right biased). Inferring the current block correctly is essential for 0% contrast trials, since they are rewarded probabilistically according to the current block (i.e., in a rightwards block, 80% of rightwards choices on a 0% contrast will be rewarded). **(c)** Mouse performance (as fraction rewarded) on 0% contrast trials, averaged around all block switches (left to right or right to left, errorbars indicate ± 1 standard error of the mean, N ranges from 38 to 141). We can see that the animals perform at a relatively stable level of around 70% accuracy, far above chance, towards the end of blocks. This drops right after the block switch to 40%, as the subjective state estimate temporarily points in the wrong direction, before recovering over the course of around 10 trials. **(d)** In the simple model, there is one static psychometric function (PMF) which maps contrasts onto logits (with the difference between leftwards and rightwards logit serving as a summary here), and **(e)** one history bias, which sums over past choices (with a leftwards actions encoded as a dot at -1 and rightwards at 1), decaying exponentially via a scalar forgetting or decay rate.

Each session involved a first 90 trials in which left and right stimuli were equiprobable, and then left and right biased blocks lasting between 20-100 trials (following a truncated geometric distribution) during which a stimulus was 80% likely to appear on one side versus 20% on the other, see **Fig. 1b**. The biased blocks define the latent state of the environment that had to be inferred by the mice, and then combined with the perceptual data – the paradigmatic form of cognition in the task. To highlight the general nature of this sort of belief formation, we will refer to the mice’s latent state estimate as their informational state. Mice were able to perform a version of this inference (Findling et al., 2023), as evidenced by their performance on 0% contrast trials (in which there is no stimulus, and so only the informational state), see **Fig. 1c**. They were on average 70% and 40% correct respectively, before and after block switches, reflecting the influence of the state. ^1^

## Uncovering behavioural details through hybrid networks

Optimal behaviour on this task would integrate signal detection theory (Green, Swets, et al., 1966) for perceptual inference (in the form of a psychometric function, PMF; **Fig. 1d**), with Bayesian inference for the informational state (Rabiner & Juang, 1986), specifically via change point detection. This history inference should be based on the sides on which successive stimuli were presented (as inferred from reward receipt or omission on each trial). Logits coming separately from perceptual inference and the updated informational state would be added for left and right choices to determine the probability of the turning direction. However, as in most cognitive tasks, there are heuristic short-cuts that subjects can exploit to perform inference more cheaply. Accordingly, and following an extensive theory-driven modelling exercise, Findling et al. (2023) interpreted the mice as making two critical approximations to estimate a subjective informational state: using their own previous choices as evidence (rather than the side on which stimuli appear, which can be inferred via the deterministic feedback); and entering these past choices into an exponential low-pass filter (**Fig. 1e**), as a form of perseveration (Gershman, 2020; Thorndike, 1911), rather than performing model-based change point detection. This strategy turns out to be only slightly suboptimal. This is because of the autocorrelation in contrast sides implied by the biased blocks, the good performance of the mice on high contrast stimuli, and the mice’s apparently appropriate choices of decay rate for the low-pass filter.

Although the Findling et al. (2023) model (which we refer to as ‘base’) is impressively competent, it is still rather restrictive, both in the type of information that it processes (only choices and contrasts) and the ways in which it processes them. We slightly generalise their model, keeping the logic the same, but allowing for a separate treatment of timeout choices (which the original model was not designed to handle), to obtain the ground floor of our model family, model ‘abcd’^2^ (details in Methods). This model predicts mice even better, but is mostly restricted in the same ways as the base model. Thus, both leave headroom for improvement that would point to cognitive components that had been omitted. We therefore took advantage of the extensive IBL dataset and determined a form of accuracy ceiling by fitting an unrestricted recurrent neural network (LSTM, but we refer to it simply as the RNN, to distinguish it from the LSTM we later use in the abcd model family; Hochreiter & Schmidhuber, 1997) to the choices of the mice. Importantly, the goal was to mimic mouse behaviour, as opposed to performing the task itself well. The RNN significantly out-performed the original model (compare abcd on the far left to RNN on the far right of Figure 2a).

**Fig. 2.**
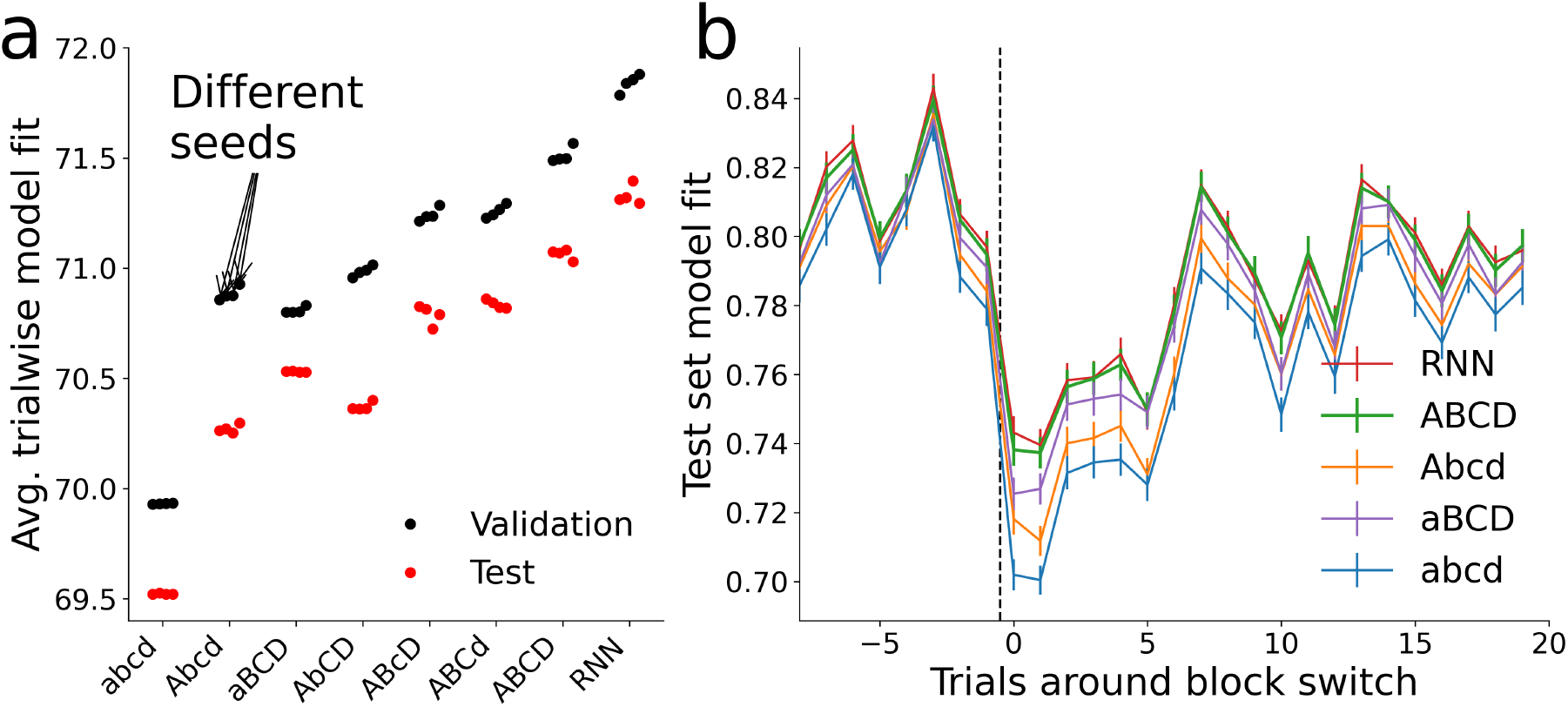
Model comparison of relevant architectures (the capitalisation of letters in the model name ‘abcd’ indicate the presence of corresponding model extensions, details below) **(a)** We show the mean log-likelihood over all trials of the validation (black) and test (red) sets as percentages, for each of four seeds with the same optimised hyperparameters. Performance on the test set is worse across the board, suggesting that it happened to include sessions which were overall more difficult to predict. Models are roughly sorted by increasing flexibility and predictive capability. (We show a complete version of this comparison in Fig. 6). **(b)**: Model fit (arithmetic mean probability assigned to chosen action) specifically around block switches in the test set. The fit quality of all models to the behaviour of the mice drops and then recovers after block switch in a manner similar to the performance of the mice themselves (Fig. 1c, errorbars show 1 SEM, N=678 for each trial).

The most revealing trials for the cognitive processes which underlie belief updating are those just after a block switch, when the mice infer that there has been a change. **Fig. 2b** shows the average fit of both the RNN and abcd to mouse choices in these periods. The RNN predicts behaviour better throughout (as can be seen by the difference in performance before the block switch), but with an even greater relative advantage right after the switch. We show later that this deviation arises because the mice track the block in a more sophisticated manner than abcd would suggest. The overall decrease in model performance after a switch is a consequence of mouse behaviour becoming noisier, as the conflict between current contrast information and recent history leads to overall closer to uniform response probabilities (see Fig. 1). With the underlying process increasing in entropy, the models necessarily become worse. However, the performance of the more competent models declines much less.

Although the unrestricted RNN fits the behaviour of the mice much better than abcd, it does not, by itself, tell us what cognitive components abcd is missing or over-simplifying. We therefore followed Eckstein et al. (2024) in augmenting the separate components of abcd with flexible feedforward (multi-layer perceptron, MLP; Rumelhart, Hinton, & Williams, 1986) and recurrent (long short-term memory, LSTM; Hochreiter & Schmidhuber, 1997) neural networks, using the performance of the unrestricted RNN as an approximate noise ceiling. We then distilled interpretable lessons from these flexible networks to enhance our simpler, cognitive, models.

The fact that the abcd model family involves two partially separate streams of information makes the first part of this procedure straightforward. We can separately augment the mapping of the contrast strength on the current trial (via a ‘PMF network’) and of a summary of past trials (via a ‘decay network’, generating the subjective informational state) onto logits, which are then added and passed through a softmax function to obtain choice probabilities. These probabilities include the third, rare, time-out option that was not included in Findling et al. 2023, the dedicated treatment of which is the major distinction between the base model and abcd. In this framework, the only limits to the predictive performance of the models are the amount of training data, the capacity of the chosen architecture, the suitability of the training procedure, and any inherent noise. Through a combination of limiting the power of the augmenting networks and distillation, we found four distinct and interpretable extensions to the base model, which, in combination, achieve predictive performance close to the full RNN (labelled A, B, C, and D, shown in figures 2a;b). In the supplement, we detail an exhaustive collection of neural network models; here, we first describe the final extensions and then show their combined effect in generating excellent performance.

The first of these extensions (‘A’, for adaptive PMF) targets the processing of contrast. It had previously been observed by Ashwood et al. (2022) that the engagement of the mice in the task (there, in the first 90 unbiased trials) might fluctuate, leading to flatter or sharper psychometric functions (Wichmann & Hill, 2001). However, that model involves punctate switching between distinct states of engagement; a flexible neural network could capture the effect of this more continuously, if appropriate. Furthermore, making this network recurrent would allow it to infer the engagement state from performance on recent past trials. Thus, we introduced an LSTM which runs in parallel to the rest of the model and provides a summarising scalar to the PMF network of our model, modulating its processing. Since it receives all information of past trials as input, an LSTM could solve virtually the entire prediction problem by itself (rather like our unrestricted RNN). Thus, we limit it to the problem of characterising sensory acuity by carefully restricting its inputs and outputs: (i) It does not receive the current contrast.(ii)It only outputs a single ‘contrast scalar’, which is input to the PMF network (a simple MLP) in addition to the current contrast. In turn, the PMF network output is forced to be a symmetric function of contrast (we restrict most of the networks to be symmetric, for the purpose of interpretability; see the section on Mirror-equivariant networks in the Methods). Thus, the LSTM cannot implement arbitrary biases, but only set the slope of the PMF. (iii) While not strictly limiting the LSTM, in the final model we regularise the change in the PMF caused by the contrast scalar, to reduce its fluctuation to the necessary minimum.

Low/high values of the scalar are nicely associated with low/high discriminabilities, as seen in **Fig. 3a**. This shows the effect of the contrast scalar by showing the output of the PMF network for a range of values of the contrast scalar (with the inset showing the marginal distribution of the scalar across trials and mice in the full model). For visualisation, we also show these PMFs in probability space directly in Fig. B7. The PMF associated with the mode of the distribution of the contrast scalar is very close to the fixed PMF of abcd (shown in Fig. B7), though it is slightly sharper, revealing that the fixed PMF is skewed by the long tail of poorer contrast sensitivity.

**Fig. 3.**
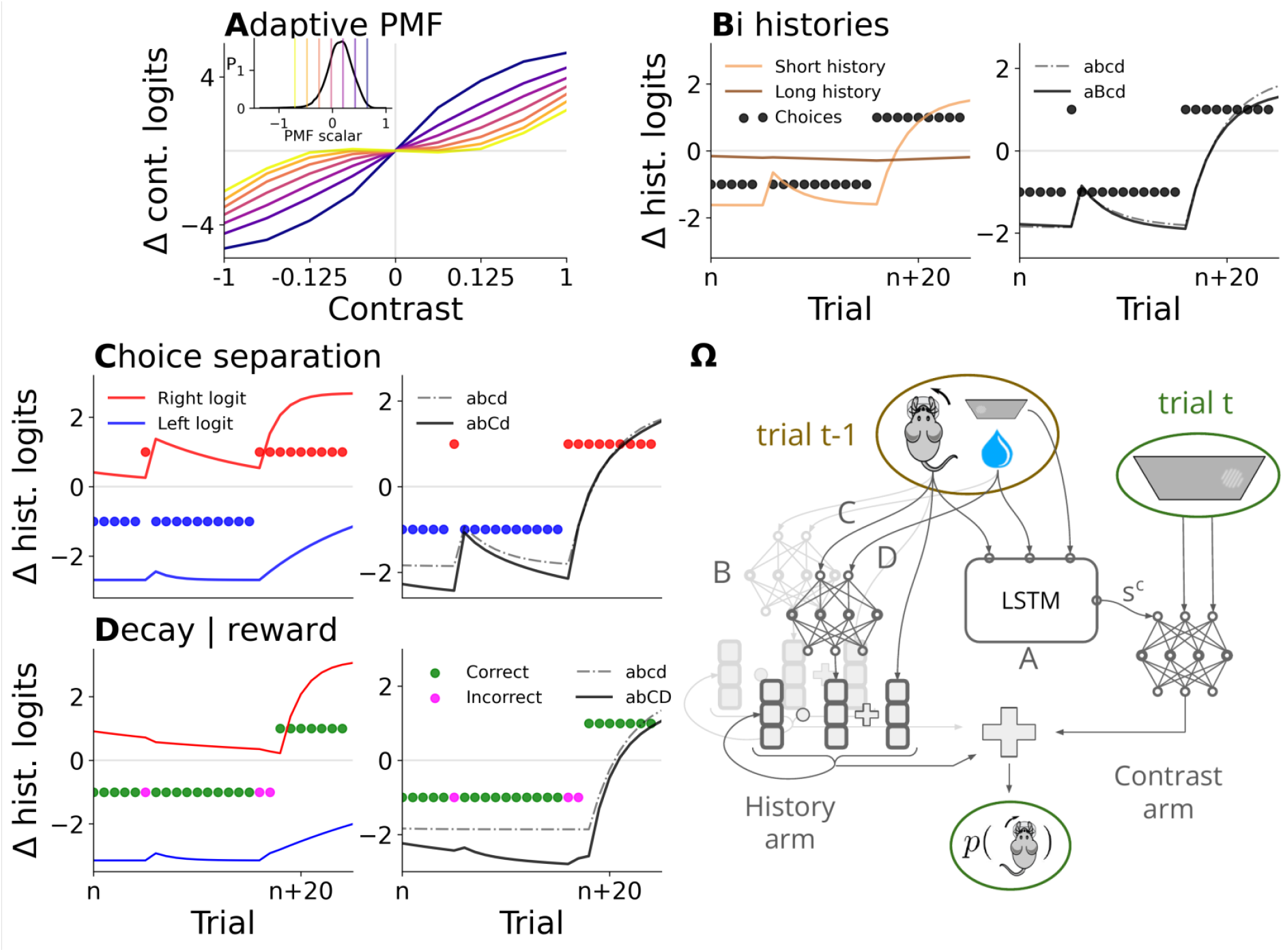
Visual depiction of the four augmentations, using an artificial trial sequence for emphasis. Together, these extensions lead to high-level, interpretable performance. The panel lettering aligns with our model naming scheme: **(A)** We make the PMF adaptive (Δ logit refers to the difference between rightwards and leftwards contrast logits), by giving it an additional input, provided by an LSTM, which determines discriminability (through a bespoke code, arranged between the networks during training). The inset shows the marginal distribution over this scalar across all sessions, vertical lines indicate different points along this distribution, which lead to different PMFs (shown in their respective colours). **(B), Left:** We use two separate exponential filters over previous actions. One (light brown) has a larger impact on the overall history component, and a shorter trial-constant. The second (dark brown) has a smaller impact, and a longer trial-constant, integrating actions (circles) over a longer time frame. **Right:** The effect of this additional filter on the overall history term. Adding this component by itself leads to a slower convergence after a drastic change in choice behaviour (seen on the very right of the plot). **(C), Left:** We track left and rightwards histories separately (blue and red line), and decay them separately via a decay vector, rather than a decay scalar. The overall history component is the sum of them. **Right:** Resultingly, the model can afford to converge to more extreme biases, given persistent choices, but also change its bias just as fast as model abcd (seen on the outlier choice at *n* + 5). It achieves this by maintaining a stronger memory of the unchosen side (weak unchosen decay). **(D), Left:** In a component that only confers benefit if combined with component C, the decay rate becomes reward dependent. The incorrect choices (fuchsia) on the incongruent trial (*n* + 5) and just after the “block switch” (*n* + 11) lead to a weakening in bias for abCD. **Right:** While the bias decreases for abCD, it does not do so in abcd, which ignores reward information. **(Ω)** Visualisation of the full ABCD model. Different networks get access to different information from the current or previous trial. Letters in the diagram denote where the different components come in: A is the LSTM which modulates current contrast processing, C gives the decay vector access to the previous choice, D gives access to the previous reward, and B provides another independent replica of this history processing (visualised in fainter grey).

The other three additions concern the processing of the informational state, and have been distilled to be directly interpretable. First, rather than just a single exponential filter over past choices (with a trial-based forgetting constant that we call the decay scalar), our network investigations suggested that, consistent with prominent findings about adaptation and reversals in other literatures (e.g., Smith, Ghazizadeh, & Shadmehr, 2006), the mice might employ more than one filter. This would allow different temporal profiles of influence from the past. We found evidence for two (component ‘B’ for binary history): a stronger one that accumulates only over recent trials; and a weaker one that accumulates over many more trials. The left panel of **Fig. 3b** shows the evolution of the two components (light brown: faster; dark brown: slower) in a set of trials manufactured to illustrate the various effects. The right panel compares the net influence on the history logits (solid) to that from the original abcd model (dashed). The main effect of the slow component is to temper the strength of inference about the block. This would permit faster inference at subsequent block switches (as in the ‘savings’ of Smith et al. 2006). The second addition to the subjective informational state involves allowing differences between the decay of chosen and unchosen actions (Funamizu, Ito, Doya, Kanzaki, & Takahashi, 2012). To do this, we explicitly split the leftwards and rightwards components of the history logits and decay them separately (in component ‘C’ for choice separation). In abcd, a single decay scalar affects all aspects of memory equally. Component ‘C’ employs a vector of decay rates, one for each logit dimension (for this it is important to realise how one can implement an exponential filter as an online low pass filter, with a momentary decay rate, we provide details in the Methods when introducing the base model). However, we again constrain this to make it symmetrical, so a leftwards action decays the leftwards history logits in the same way that a rightwards action decays the rightwards history logits (and vice versa, see Methods). Effectively, the model always uses a weaker decay factor for the unchosen action (we call a decay factor closer to 1 ‘weak’, i.e. logits experience little change), and a strong decay factor for the chosen action (respectively, we denote a decay factor closer to 0 as ‘strong’, logit memory decays more). Of course, the chosen action logit gets increased as well by the addition of that very chosen action, so these two effects interact with one another. The left panel of **Fig. 3c** shows the ultimate effect of this, with unchosen action logits (action identity is shown by the colour of the dots) decreasing very slowly. The right panel of Fig. 3c shows that, by itself, one net effect of this augmentation is to allow for faster reversals: Note how the model using component C, abCd, converges to more extreme biases than abcd, but it still reaches roughly the same level of bias as abcd after a single incongruent choice (trial n+5), traversing a larger distance.

Finally, one of the most surprising aspects of the base model that was investigated thoroughly by Findling et al. (2023), is that the subjective informational state appeared not to be affected by whether or not the choice on a trial was rewarded. This is surprising, since we might intuitively expect feedback to play a salient role. It is also suboptimal, since feedback was deterministic and informative of the true stimulus side, which relates much more directly to the current block than the mouse’s own choices (especially on weak contrasts). By distilling our best network, we found that, in fact, the decay vectors of component C *are* modulated by the outcome of the previous trial (component ‘D’ for decay according to reward). In particular, an unrewarded choice leads to a stronger decay of that chosen action’s history logit (whereas the unchosen action gets decayed almost equally in both cases). This interaction between choice identity and reward might explain why previous models did not uncover this component. The stronger decay following an unrewarded choice has a stronger effect on extreme biases, which are therefore particularly reward-sensitive, and overall, it enables a faster bias reversal in response to unrewarded choices. The left panel of **Fig. 3d** shows the operation of this component by depicting the effect of an incorrect choice on the incongruent trial (n+5). Here, an incorrect leftwards choice leads to a lowering of the leftwards logit, due to the stronger decay rate for a chosen, unrewarded action, compared to a chosen and rewarded action. In particular, the decay is so strong that it overpowers the addition (+1) of the most recent action into memory. The right panel of Fig. 3d shows that this also decreases the overall leftwards bias, because while the rightwards logit drops, it drops less than the leftwards one. In the full model, reward sensitivity thus enables faster bias switches, by reducing existing biases more quickly when they lead to unrewarded choices.

These history processing extensions combine multiplicatively, i.e. in the full model there are four (two histories times two rewards) **vectors** of decay (splitting leftwards and rightwards history)

### Behavioural fits

We show the validation and test performances of the relevant networks in **Fig. 2a** (supplementary figure 6 includes a complete comparison, including the network models that were distilled to create these components). The final interpretable architecture, ABCD (remembering that letter capitalisation indicates the presence of an augmentation), gains almost all the available increase in performance over the original base model to the unrestricted RNN. Confirming this excellent performance, **Fig. 2b** shows that ABCD captures closely the improved fit of the RNN around the time of block switches.

The contrast scalar, capturing sensory acuity, plays a critical role in this improved performance, as seen by the performance of Abcd in **Fig. 2a**. Though notably, it leads to a somewhat larger gap between test and validation performance than models without A exhibit, indicating that it does not generalise flawlessly to those never learned-on sessions. We dive into the workings of this component, in the context of ABCD, in **Fig. 4**. First, **Fig. 4a** shows the calibration of the contrast scalar by comparing empirical (green dashed) and predicted (red) PMFs, averaged over trials on which the contrast scalar is in each of four quartiles. The fit is excellent – far better than the fixed PMF of abcd. Thus, the contrast scalar accurately reflects the perceptual sensitivity of the animals, and relates directly to the quality of behaviour. This allows it to capture the momentary motivational state of a mouse as the session evolves. One would thus expect that its overall statistics would covary with performance. Consistent with this, **Fig. 4b** shows the marginal densities of the contrast scalar across all trials of each of the 403 sessions, sorted by the overall reward rate of the sessions (left line). Sessions with higher reward rates have higher (and less variable) values of the contrast scalar.

**Fig. 4.**
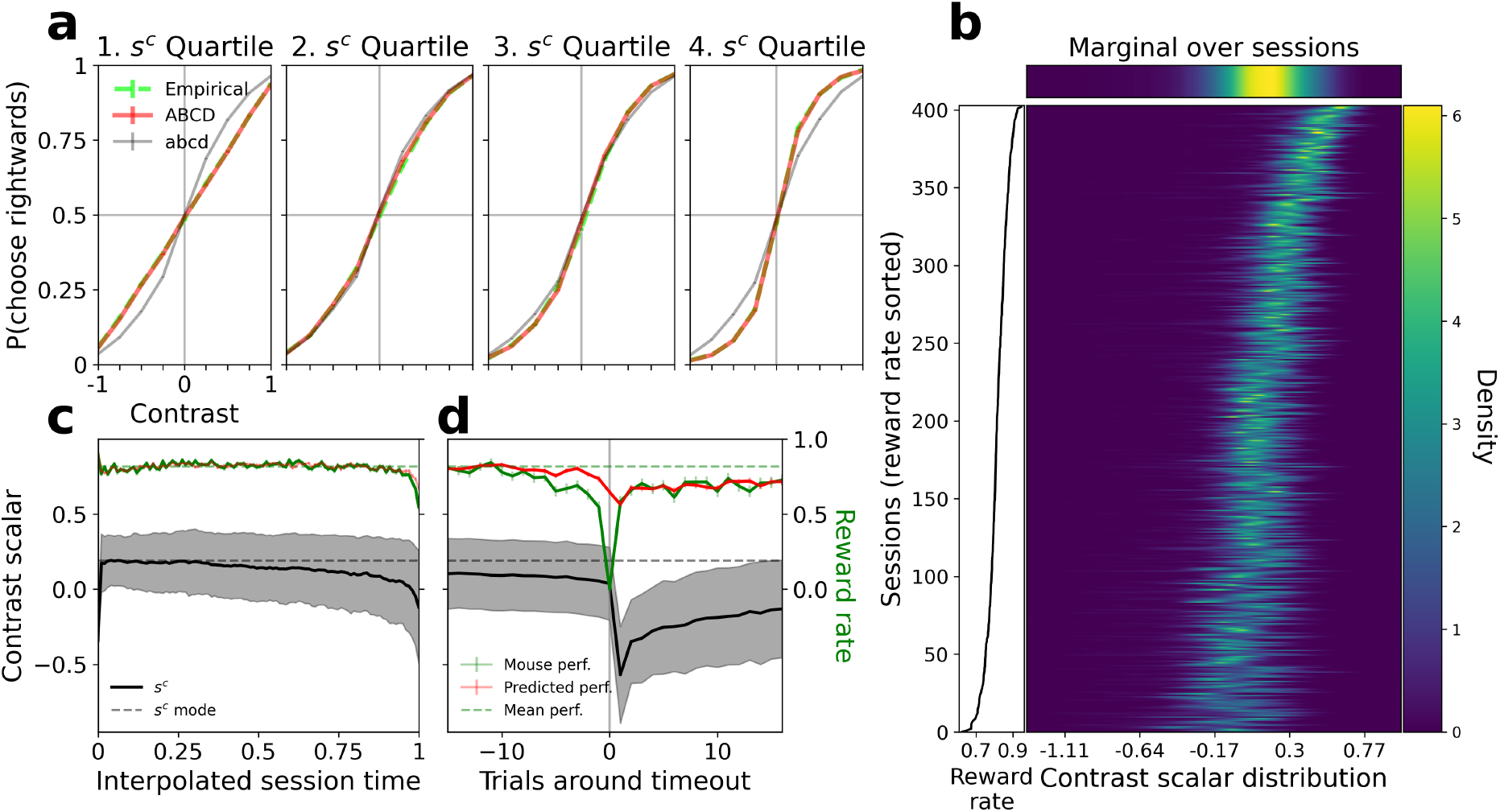
Contrast scalar values reflect the behavioural engagement of mice on a trial-by-trial basis. **(a)** Calibration of the predicted PMFs of ABCD and abcd versus the empirical PMFs, across a quartile split of the trials based on the contrast scalar *s*^*c*^ (errorbars are small, but depict ±1 SEM, each datapoint covers at least N=6692 trials). The effect of the scalar upon ABCD is obviously very well calibrated, as reflected by the close agreement between the predicted and empirical PMF. abcd is much less flexible (though it does have some adaptability due to the perseveration component), and thus does not accurately reflect behaviour for the extreme quartiles. **(b)** Relationship between the contrast scalar of ABCD and the reward rate across sessions. The heatmap shows the marginal distribution of the scalar over all 403 training sessions, sorted by their reward rate, decreasing from top to bottom (the plot on the left indicates the reward rate). Higher values of the contrast scalar are associated with higher reward rates. **(c)** Averaged and interpolated evolution of the contrast scalar across sessions (we linearly map each session of whatever length onto the range from 0 to 1). Errorbars show ±1 SEM, whereas the shaded region indicates ±1 standard deviation to give a sense of spread, N=403 training sessions. The scalar has a strong tendency to decrease over a session, along with reduced task performance, presumably due to disengagement. **(d)** Averaged contrast scalar around each first timeout choice (out of 403 sessions in the training set, 194 have a timeout choice), along with actual and predicted reward rates. Errorbars and shaded region follow the same logic as in (c). The lowest number of trials for a data point in this plot is N=151, due to some sessions ending soon after the first timeout. A timeout choice leads to an extremely marked decrease in the scalar, which then, on average, slowly decays. As shown by the green line, the reward rate of the animals decreases before a timeout, which is not fully captured by the contrast scalar.

This relationship between the contrast scalar and performance also holds during the course of the sessions, both broadly and minutely. **Fig. 4c** depicts the average value of the contrast scalar per trial across sessions (warping each session to be the same notional length of 1). The steady decrease in the contrast scalar is consistent with the progressive disengagement of animals from the task and the drop in overall performance. More finely, one of the major signatures of disengagement are timeouts, when the subject does not respond on a trial for 60 s. **Fig. 4d** shows the results of averaging the contrast scalar around the first timeout in a session (for sessions that contain at least one). The drop after the timeout is consistent with the degradation in performance and the empirically observed level of accuracy is well reflected by the predicted reward rate, enabled by the lower contrast scalar. However, the contrast scalar does not allow for the accurate prediction of a timeout choice, nor does it fully capture the small lead-up in the form of a lowered reward rate. It implements a reactionary description of engagement, and acts as a low-pass filter, integrating performance over a longer time horizon.

The other interpretable components of the model work together to improve the subjective informational state inference. **Fig. 5** and Fig. B12 show different aspects of this. First, **Fig. 5a and b** show the decay vectors for the faster and slower history filters, for chosen and unchosen actions and depending on whether they were rewarded or not. For the dominant, faster filter, the unchosen action decays much less than the chosen action; this effect is exacerbated if reward is not provided. For the slower filter, the pattern is more complicated, with an unrewarded unchosen action being suppressed rather than boosted. Overall, the decay rates of the slow filter are very weak, leading to large integration windows. As previously hinted, the full model enables a stronger decay of the chosen actions’ memory in response to reward omission, leading to a relatively weaker increase in bias (or even a total weakening of bias). We visualise this in **Fig. 5c**, which shows histograms of the total change in the subjective informational state for abcd and ABCD as a function of the starting value for various choices. ABCD is more expressive than abcd: the simple model treats the history-based bias as a 1D quantity (all updates lie on a line, by design), leftwards and rightwards biases trade off directly with each other. For ABCD, however, the model has an additional degree of freedom in its bias, allowing it to slightly modify how amenable the history term is to future changes. Quite saliently, at the extreme ends of the subjective informational state (large positive starting logit difference), a belief-congruent, but unrewarded, choice, leads to a lowering of this belief, something which cannot be achieved by abcd. The presence or absence of reward thus does affect the animal’s belief updates.

**Fig. 5.**
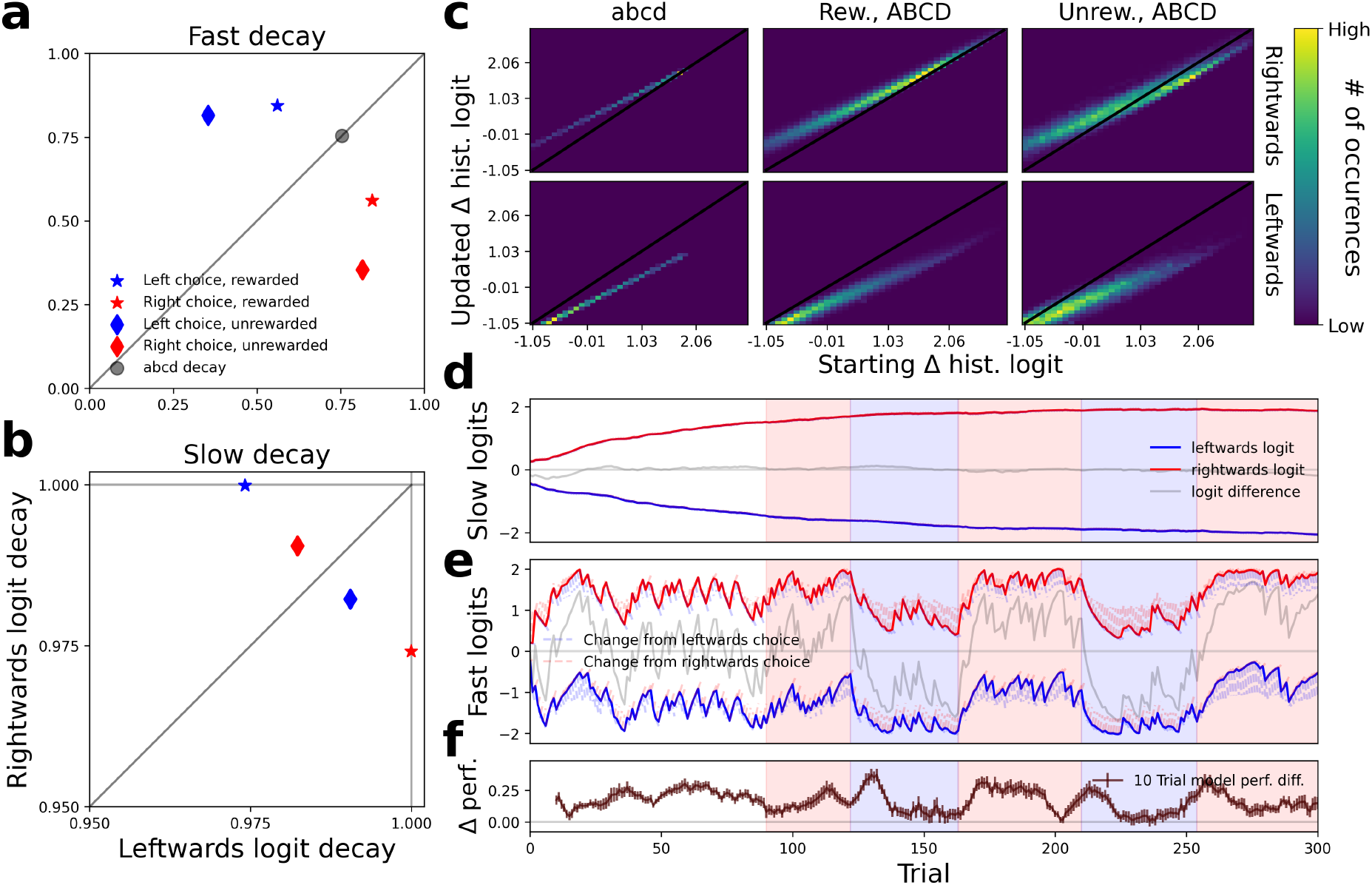
Informational state inference becomes reward-sensitive and overall more sophisticated **(a)** Decay factors of the fast history (larger weight), for the chosen and unchosen history logit, depending on whether the choice was rewarded. The grey circle denotes the single, reward-independent decay factor of the abcd model. **(b)** Decay factors of the slow history (smaller weight). Note the reduced axes range, due to the weak decay factors. **(c)** Heatmaps of the resulting history bias (i.e. logit difference), given different types of past trial (each heatmap’s colour scheme is normalised individually). We focus on the positive logit differences (i.e. rightwards biases), as these changes are symmetric. Due to the coupling of left and right logits in abcd, the distribution of changes is constrained to a line. Additionally, they are independent of reward. Model ABCD is more expressive and can create the same logit difference through a range of values. The heatmap of rewarded choices appears mostly similar to abcd, but unrewarded choices differ notably. In particular, given a large initial difference, an unrewarded rightwards choice leads to a lowering of this bias (top right quadrant of top right heatmap). **(d-f)** We now consider a specific session type (1) and average over all its training sessions (N=74): **(d)** History component of the slow history. It is a subtle effect, but the slow history tends to counteract the current block (the grey difference line runs counter to the block). **(e)** The fast history component of the overall history bias of ABCD. The whiskers indicate how the logits would evolve depending on the choice. **(f)** Mean performance difference between models (difference between probabilities assigned to the chosen action by ABCD and abcd), summed over the last 10 trials (errorbars denote ±1 standard error of the mean). ABCD always predicts better, particularly after block changes (see background colour). We show the last three panels for other session types in the Supplement.

The remaining plots in **Fig. 5d-f** show a birds-eye view of the workings of the informational state in ABCD by averaging over all the sessions of the most prevalent type (type 1). **Fig. 5d** shows the evolution of the left and right logits for the slow filter, together with their difference. Interestingly, the decay implements dynamics which imply that the slow filter often slightly counteracts the current block appropriate choice (see in particular **Fig. B9**, depicting a session type in which rightwards blocks dominate and the slow filter implements a consistent leftwards bias). This contributes to the model’s ability to implement faster reversal behaivour. The initial values for both filters are learned along with the other components of the model – and the initial bias (perhaps because of the prevalence of some session types) and the slow rise (perhaps because of the initial 90 unbiased trials) was unexpected.

Similarly, **Fig. 5d** shows the evolution of the left and right logits for the fast filter, together with their difference and the changes that would have been made according to leftwards or rightwards choices. As we already saw in general in **Fig. 5c**, whenever the logits take on very high (or, in fact, very low) values, they become amenable to the greatest change.

Taken together, these additions lead to notably better predictive performance by ABCD, particularly around the critical time of the block switches. We see an overview of this performance difference in **Fig. 5e**, averaged across a small causal window of 10 trials. While the differences are rather spread out, the block switches (and other times at which the bias changes quickly – evident from the fast filter) often lead to a greater advantage for the full model. We prove the significance of this quantitatively in Fig. B11. Part of this is ABCD’s ability to capture faster bias reversal, which can be seen very clearly in Fig. **B12a**. This depicts the trials in which ABCD holds the greatest advantage over abcd for that particular session type. By chance, the histories of the two models start out at very similar values before the block switch, but we can see that the average history of ABCD drops faster, while providing a better explanation of mouse choices.

## Discussion

We presented the results of adapting and applying a systematic methodology to understand a fundamental cognitive operation that is common in a wealth of paradigms. In this framework (Eckstein et al., 2024), we start from a cognitively penetrable model (Findling et al., 2023; Wilson & Collins, 2019), augment its components with flexible neural networks until getting suitably near a noise ceiling in performance, which is also estimated by an unrestricted neural network (Dezfouli, Griffiths, et al., 2019), and then analyse the result for generalisable lessons. Doing this effectively is a form of meta-learning (of the population of subjects rather than a population of tasks; Binz et al. 2024) and requires substantial data - for which we exploited the IBL (The International Brain Laboratory et al., 2021). While the effect size of the improvement in model fit was modest, it represents a clear improvement over existing models and undoubtedly captures behaviour more holistically, an important property for theoretical understanding and further analyses.

In particular, we find that the sensory acuity of the mice strongly fluctuates during the course of a session. On the belief estimation side of the task, it turns out that mice apply subtle tweaks to the simple exponential filter model: they use two separate filters, one fast and one slow, they track left- and rightwards biases somewhat separately, and their belief updates are, in fact, reward sensitive. Interestingly, approaching the performance ceiling on this task did not require accommodating inter-individual differences beyond the fluctuations in sensory acuity (which appear to be dominated in large part by changes within an individual during a session). Belief estimation through components BCD was able to describe well the entire population, as, if there would have been notable differences, the RNN would have presumably picked up on them and adapted during the course of its predictions, leading to a more notable predictive advantage of this model. Similarly, components BCD did not need access to the current motivational level of the animal (contrast scalar *s*^*c*^) to perform well, though this might be an interesting area for future extensions.

It is an important next step to follow (Findling et al., 2023) and use the electrophysiological data in The International Brain Laboratory et al. (2023) to test for the existence of the four augmentations we found. The contrast scalar motivational signal might covary with the activity of neuromodulatory arousal systems (McCormick, Nestvogel, & He, 2020); it would then be compelling to attempt to interpret the way that the LSTM that generates this signal does so by exploiting past errors to infer motivational state. The implementation of the putative multiple timescales of the decay is more obscure (Cavanagh, Hunt, & Kennerley, 2020), although Findling et al. (2023)’s findings of regions with activity covarying with the decay of abcd would be an excellent starting point. Neural data are important to mitigate the possibilities of model mimicry.

The existence of substantial datasets such as IBL is encouraging active exploration of various approaches to improving on compact models, including the direct or indirect use of small networks (Ji-An, Benna, & Mattar, 2023; Schaeffer, Khona, Meshulam, Laboratory, & Fiete, 2020), forms of regularisation (Dezfouli, Ashtiani, et al., 2019; Miller, Eckstein, Botvinick, & Kurth-Nelson, 2024) and even evolutionary optimisation for program induction (Castro et al., 2025). The last of these also noted the great complexity devoted to the equivalent of the informational state, with sophisticated forms of perseveration (Gershman, 2020; Thorndike, 1911). For all these methods, as with ours, fitting is straight-forward and automatic, deriving the understanding is more bespoke. In our case, the particular danger is that flexible networks, such as the LSTM that generated the contrast scalar, can solve the entire problem themselves in ways that are non-transparent; we suffered exactly this problem when we allowed the decay vectors controlling the informational state to be influenced by a similar scalar. This difficulty may help explain the remaining small gap with the performance of the unrestricted RNN, which can adjust to differences in block inference that become evident as sessions evolve.

Nevertheless, we have shown that we can improve upon an already excellent model of a cognitively important task by integrating data-driven and theory-driven methods. Mice turn out to be more competent at latent state inference than previously thought, whilst also suffering from continuously changing levels of engagement. This offers important lessons for characterisations of other cognitive tasks.

## Methods

### Further task details

Sessions ended when any of: (i) the mouse had been working for more than 90 minutes, (ii) the mouse failed to do more than 400 trials in the first 45 minutes (in which case it performed so poorly that we anyhow excluded the session from our analysis; see section on Network training and evaluation, or (iii) the mouse had completed over 400 trials and the median response time (defined as the time from stimulus onset to the registration of the response) over the past 20 trials was over five times longer than the median response time over the previous trials in the session. The third condition was the most common reason for a session ending, with behaviour often degrading towards the end of a session as the animals disengaged, possibly due to satiation.

Mice performed varying numbers of sessions while being recorded, ranging from just one up to sixteen, with a median of three. For the purpose of reproducibility, the IBL employed the exact same sessions for all mice while they were recorded (the training sessions, on the other hand, were created randomly on the spot). By “the exact same”, we mean that the contrasts which were presented and the choices which were rewarded followed a fixed sequence, so we call these prototypical sessions “session types”. There were twelve different session types, which were presented to each mouse in the same order (though some mice, presumably accidentally, were presented the same session types twice). As a consequence, the early session types were considerably more frequent in our data set than later ones. The count of session types in order is: 107, 98, 96, 90, 32, 30, 30, 24, 13, 8, 6, 5. The prevalence of the first four session types led to visible trends in some of our performance plots. For example, since there were only so many block switches to observe, if the contrast on the third trial after a switch was, by chance, usually a strong one, the predictions on this trial will be relatively better than surrounding trials, as seen in **Fig. 3b**.

### Hybrid network ladder

We introduced flexibility progressively, starting from model abcd (which roughly generalises Findling et al., 2023), defining what might be called a ladder of hybrid neural networks (Eckstein et al., 2024). abcd, which is at the bottom of the ladder, is itself a generalisation of a classical, simple, and interpretable exponential filter in a multinomial logistic regression framework (covering three possible responses, rather than just two), which we call ‘base’. The neural network architectures are designed as explicit extensions of this model, enjoying two structural forms of added flexibility and complexity. We begin by detailing the simplest model and then introduce abcd and the other relevant neural network extensions.

#### Level 0: base

Our simplest model, abcd, is a generalisation of a model that employs the same general framework as Roy, Bak, Akrami, Brody, and Pillow (2021) and Ashwood et al. (2022), namely logistic regression with selected features. This model has been widely applied to IBL data (and is comparable to the model of Findling et al. (2023)). To acknowledge its foundational nature, we call this model ‘base’, but do not consider it part of the ladder, due to its naive handling of timeouts. For the feature based on the current stimulus, we produce contrast logits by applying a tanh transform (as suggested by Roy et al. 2021) elementwise to the contrasts **c**_*t*_ on trial *t*. **c**_*t*_ is a two-dimensional vector encoding the strength of the contrast on the left and the right side of the screen (at most one of which will be non-zero). Since we also consider timeout choices, we apply a multinomial logistic regression framework. We produce three logits to turn into probabilities over three possible choices. The tanh is thus followed by a learnable linear transformation comprising a matrix ℳ_*c*_ ∈ ℝ^3,2^ and a bias **b**_*c*_ ∈ ℝ^3^:

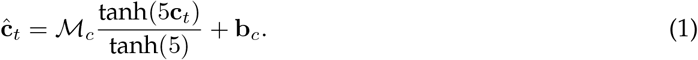

We use ^ to indicate that raw information has been processed into a logit representation associated with the action propensities for the different choices.

The contrast logits are supplemented by history logits, which summarise the recent history of choices as an informational state, and act as a prior for the current choice. This can reflect largely appropriate considerations about the current biased block, but also other perseverative tendencies of the animals. An action is represented as a three-dimensional one-hot vector. The first entry encodes whether a leftwards choice occurred, the second a timeout “choice”, and the last dimension indicates a rightwards choice. After a trial, the current informational state or memory over previous choices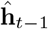is updated by multiplying it with a scalar decay constant 0 *< d <* 1, and adding the most recent action **a**_*t*−1_ into memory:

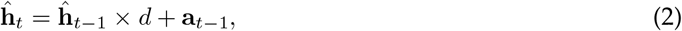

with the starting point ĥ_0_ being fitted during training. This repeated multiplication implements an exponential decay of the influence of past choices. Note that this is the same computation as a convolution of choices with an exponential filter

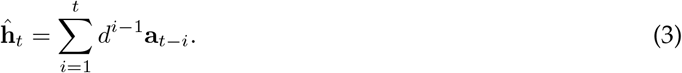

However, thinking of it as a trialwise update will later allow us to modulate the momentary (trialwise) decay rate *d* of the filter, which will prove essential.

The ultimate choice probabilities **p**_*t*_ are generated by adding the two components (weighting the history by a learnable scalar *w*) and passing them through a softmax function:

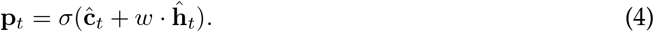

With the softmax defined as

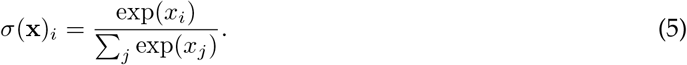

This is the most basic approach to modelling our behaviour of interest. Note, however, that this model was not designed with timeout ‘choices’ in mind, which were simply ignored in previous work. We directly extend the logistic regression to cover all three possible responses, but do not treat timeouts any differently within the decay equations. Since timeouts are obviously special occurrences, this could be considered somewhat of a straw-man model. We address this obvious shortcoming through our next model.

#### Level 1: Model abcd

The level 1, simplest model of our family, is a modest generalisation of the level 0 model which, includes more flexible processing of contrast, and also a dedicated decay scalar for the timeout choice (but still using the same functional form as equation 2).

In particular, rather than picking a specific functional form for the mapping from contrasts to logits, we input the current contrast into a network (contrast network), which then produces a three-dimensional output vector of logits. It does so in a symmetric way: a 25% leftwards contrast, for example, produces the same logits as a 25% rightwards contrast, only flipped (see the section on Mirror-equivariant networks for details). This contrast network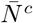 (using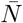 to denote the opera-tion of a mirror-equivariant feedforward multilayer perceptron, or MLP) thus performs the simple transformation:

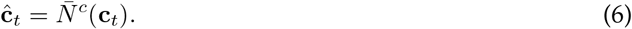

We tried various network sizes, as discussed in the section on Network training and evaluation.

To acknowledge the special role of timeout choices, we effectively employ four different decay rates to update the informational state, *d*_1_, *d*_2_, *d*_3_, and *d*_4_, like so:

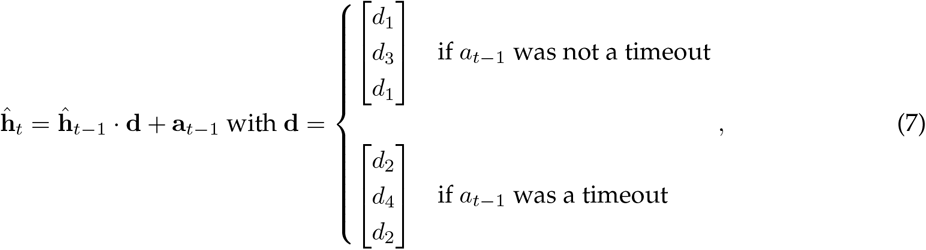

where · denotes a Hadamard product. Intuitively, leftwards and rightwards logits decay with a single decay constant most of the time (*d*_1_), but timeout logits have their own decay constant, and the model also gets to use specialised decay rates in response to a timeout (as they are unusual events).

In sum, this represents a straightforward extension of the widely-used base model to our setting: it allows for the dedicated handling of timeouts and does not restrict contrast processing to any specific functional form (though it does enforce symmetry).

#### Level 2: Flexible history processing - BCD

In the classical exponential filter, the one-hot vectors of previous actions decay by a constant proportion on each trial. This implies that memory is multiplicatively weakened through the decay constant *d*. Initially, we employed a neural network to directly map three-dimensional previous memory vectors onto three-dimensional current memory vectors, allowing for more complex patterns than the straightforward decay from one-hot towards zero. However, to reveal the interesting patterns of action-dependent memory decay, it turned out to be more beneficial instead to use a neural network to produce a vector of decays after considering events of the past trial (decay network):

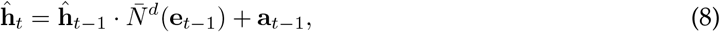

**e**_*t*−1_ denotes the events of the past trial. We can increase the capacity of the models by including more information here. Note that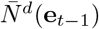takes over the role of *d* in eq. 2. For the simple abcd model, we only provide information about whether the response was a timeout or not. This effectively, as discussed, grants the model two decay vectors, but within one such vector, the decay of the leftwards and rightwards logit is always exactly equal, thanks to the symmetry constraint of the neural network. The timeout logit has an independently fit decay rate, as the central position of the decay vector is not constrained by symmetry.

Going one step further, we can pass the entire one-hot action vector to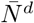, i.e. **e**_*t*−1_ = **a**_*t*−1_. This allows for three separate decay vectors, but the vectors for leftwards and rightwards choices are flipped versions of one another. This allows for separate decay of the chosen versus the unchosen logit, giving us model component C.

Alternatively, or in addition, we can include information on whether a trial was rewarded or not in **e** (note that this is passed as additional information, and does not play into the symmetry considerations of the neural network). This then doubles the number of decay vectors, allowing for separate dynamics depending on reward, model component D.

Lastly, model component B simply makes use of two separate memory processes, and two separate decay networks, modifying the update equation to this:

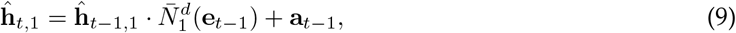

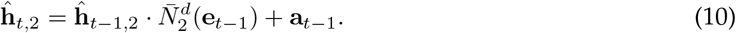

Eq. 4 now uses two learned weights to factor in the two memories separately, becoming:

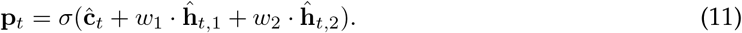

These two independent histories end up becoming the fast and slow history filters (though they are in now way pushed into these specific roles). These components can be introduced into the model independently from one another, giving us the family of models from abcd to aBCD.

We also experimented with feeding the action through an “encoding” network, before adding it to **ĥ**_*t*−1_, which allowed for the processing of e.g. previous reward and contrast, but the resulting architecture proved hard to interpret.

#### Level 2: Additional flexibility through recurrent networks - A

Yet better predictive performance came from adding an additional separate, modulatory, network that runs in parallel with these previous extensions and succinctly summarises past information (i.e. excluding the current contrast). This network provides an additional, single scalar input to the contrast network, thereby modulating it. For this we use a recurrent network (an LSTM; Hochreiter and Schmidhuber 1997), preserving its own hidden state across trials, unlike the previous components. Therefore, this network is much more powerful than the others and could, in the worst case, absorb the explanatory dynamics within its difficult-to-interpret hidden states.

We limit the power of the modulatory network in various ways: (i) We only allow it to pass a single scalar to the contrast network, reducing its expressiveness. (ii) We optionally regularise the first derivative of the effect that the LSTM output has upon the implemented PMF, using the *L*^1^ norm (see the section on Network training and evaluation for details), exerting pressure on it to capture only those fluctuations that are necessary. (iii) We use a mirror-equivariant contrast network, forcing it to treat leftwards and rightwards contrasts of the same strength as equal but opposite in how they affect the upcoming choice. This ensures that any biases are determined by the history processing (and, in particular, that the two streams do not trade off biases), and the scalar can only modulate PMF steepness (i.e. sensory acuity); for details, see section on Mirror-equivariant networks.

The introduction of this modulatory network was necessary in the first place because we found that temporary fluctuations in the behaviour of the animal (presumably through factors like motivation), and possibly individual differences across animals, are substantial components of the explainable variance. If an architecture lacked an explicit mechanism to handle them, they were captured as much possible by the other networks, rendering them needlessly complex and complicating the dynamics of their representations. Thus, while this additional network provides an extra component to interpret and an additional input within the contrast networks to understand, handling it as an explicit modulator at least makes that component transparent. Furthermore, as we showed, it is readily interpretable.

To pass the summarising contrast scalar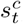, we adjust the PMF network equation as:

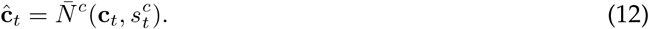

Granting us component A.

Although we ultimately distill it away, we also considered allowing the decay vector to be influenced by the output of the modulator network:

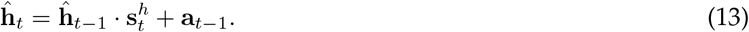

In the complete case, the contrast scalar and the decay vectors are produced by separate learned linear readouts from the hidden state of an LSTM, which receives as its input the whole list of **a**_*t*−1_, **c**_*t*−1_, and the reward *r*_*t*−1_:

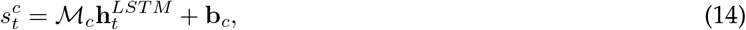

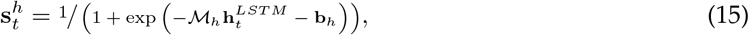

where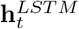 is the hidden state of the LSTM on trial *t* and we applied an elementwise sigmoid function to constrain the decay vector entries between 0 and 1. We call this auxilliary architecture A+ID, as it uses the component A of the family of models, but also lets the LSTM perform **I**nference over the **D**ecay. We briefly discuss this model in the next section. We work with a standard LSTM architecture throughout.

The highest rung on the hybrid network ladder is a completely unrestricted recurrent neural network (RNN), which consumes all input data at once and indiscriminately on every trial (**c**_*t*_, **a**_*t*−1_, and *r*_*t*−1_) and directly produces output probabilities over the three choices. In particular, this network is able to implement arbitrary interactions between the input streams. We again use an LSTM for this, but to differentiate it from the LSTM of component A, we refer to it as the unrestricted RNN throughout.

This concludes the description of all relevant architectures. The full model comparison is shown in **Fig. 6**.

**Fig. 6.**
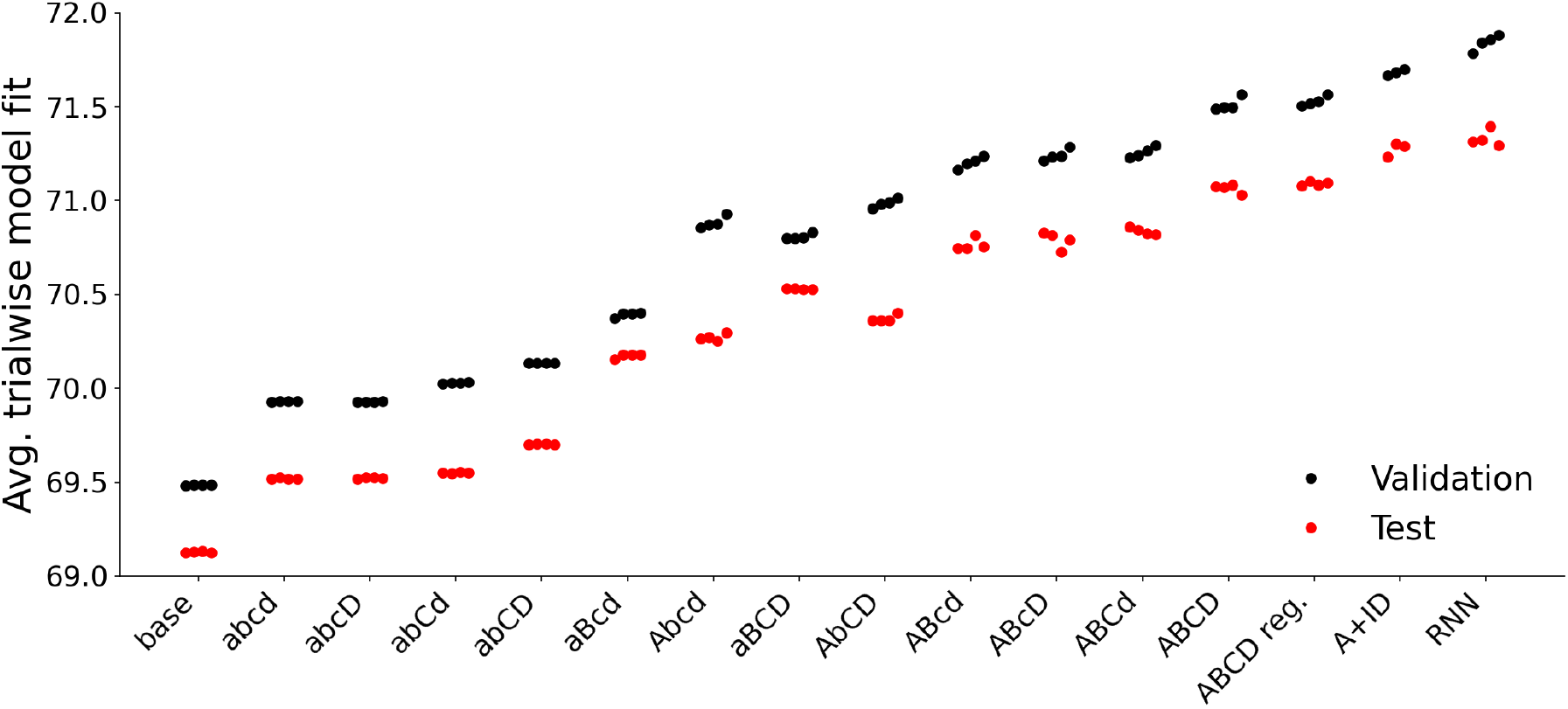
Model comparison of all architectures, showing the mean trial-wise log-likelihood on the validation (black) and test (red) sets for each of four seeds with the same optimised hyperparameters. Models are roughly sorted by increasing flexibility and predictive capability. We include a version of ABCD which was regularised on the amount of change implemented via the A component, which performs very similar to ABCD itself. The auxilliary architecture A+ID performs a bit better still than the ABCD-family.

### Model distillation

We experimented with a number of architectures, studying which additions were beneficial, and trying to distill such additions into simpler components. This ultimately succeeded through architecture A+ID, which splits contrast and history processing, our minimal assumption, but modulates both through the usage of a flexible LSTM, making them very powerful. Through this architecture, we quickly noticed that there were strong patterns in the LSTM selected decays, which we then distilled into simpler networks.

Most saliently, the LSTM always chose to apply a stronger decay to the chosen action than the unchosen one. We therefore allowed for this flexibility in our decay, extending the previous decay scalar framework (component C).

When looking at the biases which A+ID implemented through its history processing, we noted that one component (say, the leftward logit) always fluctuated much more strongly than the other (correspondingly, the rightward logit). We considered this the usage of a degree of freedom which the model had, but that was not desirable, as there was *a priori* no reason for animal behaviour to be asymmetric in this way. However, any attempts to force the model to respect symmetry failed, as it led to decreased predictive performance. Speculating that the model might need to make use of a fast and a slow changing memory, we explicitly allowed for two separate histories, introducing component B. This turned out to be the needed extension, as we were able to enforce symmetry, if only we allowed for two histories.

Lastly, we considered previous reward an interesting feature from the start and included it in most attempts at model extensions. It was therefore a natural inclusion within this framework of adaptive decay vectors and worked immediately.

Component A, which currently relies on an LSTM, is an obvious target for further distillation. We did notice during model fitting that it does not strictly need all the provided input, and in fact performs almost as well by just receiving previous choice and reward information (ignoring the previous contrast). Further studying the workings of this network is thus an interesting direction for the future.

### Network training and evaluation

We trained the models on behaviour in a total of 539 sessions of 139 fully proficient mice that were simultaneously being recorded using Neuropixel probes (these data are substantially analysed in Findling et al. (2023); The International Brain Laboratory et al. (2023)). Our sessions were selected from the total of 693 sessions in The International Brain Laboratory et al. (2023) (apart from 6 sessions that were more recently added) based on two exclusion criteria, which were determined before analysing the data, and loosely follow The International Brain Laboratory et al. (2023): we removed sessions with fewer than 400 trials (which is more stringent than The International Brain Laboratory et al. (2023)), as well as sessions with poor performance on easy contrasts, namely fewer than 90% correct responses on 100% contrast inputs. This left the 539 sessions. We did not remove sessions or trials based on reaction times or neural data properties, unlike The International Brain Laboratory et al. (2023). The rationale behind these choices was that behavioural quality was of high importance (thus the stricter limit on the number of trials, as performance needed to be poor for the automatic protocol to end a session before trial 400), but the quality of the neural data or strong temporal consistency (which eases neural analysis) was less relevant.

We trained our networks on a training data set, consisting of 403 sessions (∼75% of all sessions, in total encompassing 262,569 trials). We evaluated the fits on a validation set, comprising 68 sessions (representing ∼12.5% of all sessions, 43,278 trials). We used this to determine the best hyperparameters for our training procedure. The results we report in the paper were achieved on the final test set of the same size, which was only used once at the very end (68 sessions, 43,968 trials).

We stratified our data split according to two important criteria: session length and session type. As noted, a session usually ended when reaction times reached a level that was substantially above average; thus the end of a session was almost always marked by degraded behaviour. The length of a session was thus correlated with the extent of good quality behaviour. Therefore, we ensured that each split had roughly even proportions of all different types of session lengths. To this end, we divided sessions into quartiles based on their numbers of trials, and distributed sessions from each quartile into our train-validation-test split (following the 75%-12.5%-12.5% proportions). Behaviour had to be poor to lead to a session with less than 400 trials, which is why we excluded such sessions. To encompass the possibility that sessions with only just a few more than 400 trials may have had similarly bad behaviour to those we excluded for having less than 400 trials, we employed a separate bin for all sessions with 405 trials or fewer (of which there were 28), which we also spread out approximately uniformly across the splits. The second criterion was the session type (of the 12 canonical types mentioned above). To preclude our models from being specialised to just a subset of the presented session types, we ensured that (the already biased collection of) those we had were distributed relatively evenly across the data sets, introducing another level of stratification into our splitting scheme.

To fit the training sessions, we minimised the negative log-likelihood which a model assigned to the actions the animals actually took (in the batch and data set under consideration). That is, we minimised

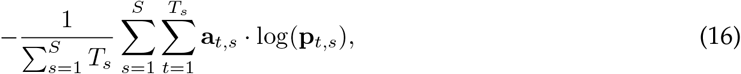

where we index the one-hot vector of the chosen action **a** and the model generated probabilities **p** by the trial *t* and session *s*. Here, *S* denotes the number of sessions used and the second sum goes over all trials *T*_*s*_ in a specific session, as sessions were of variable length. We divide by the number of trials considered, to have a properly normalised quantity.

We performed an independent hyperparameter sweep for each network architecture, to determine the ideal model capacity and training procedure. We varied the batch size of the number of sessions on which to evaluate the gradient (since we were also training recurrent networks, sessions had to be evaluated in their entirety), trying out batch sizes of 8, 16, 32, and 64 sessions. We used the Adam optimiser (Kingma & Ba, 2017), with different learning rate initialisations, considering 10^−4^, 10^−5^, 10^−6^, and regularised using different strengths of weight decay, 10^−4^, 10^−5^, 10^−6^. We also varied the number of units in the hidden layers of the networks, trying 4, 8, 16, and 32. This applied separately the contrast network and the LSTM. The overall number of values to set was too large to search exhaustively. Luckily, a few parameter settings were generally favourable across a range of network types: A learning rate of 10^−4^ clearly outperformed other settings, and the number of hidden units was generally best set at the intermediate values of 8 or 16. Similarly, networks either benefited from rather small or large batch sizes, so the intermediate batch size of 32 proved generally unhelpful. Therefore, we sometimes limited searches to the better values.

To gauge the reliability of our training procedure, we paired each hyperparameter combination with four different seeds, allowing us to compare performance across four random initialisations. As seen in **Fig. 2**, the performance of those seeds for the stronger models could differ. To ensure that models had trained to completeness (though this can be hard to determine, as steep improvements in performance may occur after many training steps without obvious improvement Power, Burda, Edwards, Babuschkin, and Misra 2022), we trained for 350,000 epochs, by which point it appeared that the networks had thoroughly converged in terms of both training and validation loss. We evaluated the loss of each network on the validation data set at least every 20 training steps, and used the best-performing model on this metric as the representative for its architecture (ignoring possible differences to worse performing seeds).

For the full ABCD model, our winning hyperparameter setting was: batch size 8, learning rate initialisation 10^−4^, weight decay 10^−4^, LSTM dimensionality 8, contrast network dimensionality 16.

The dynamic outputs of the LSTM in the A-networks could fluctuate more than necessary. We suppressed this by adding a regularisation term to the loss function, reducing the change in the outputs of the LSTM to what was necessary to describe behaviour well. We thus computed the changes within the PMF that was implemented by the contrast network, given the contrast scalars of adjacent trials (it does not suffice to punish the contrast scalar directly, as it can simply modulate the contrast network in the same way as it normally does by varying on a smaller range). We then computed the *L*^1^ norm of these difference terms, and added a weighted sum of them to our loss. We chose *L*^1^ rather than *L*^2^, so as to not punish strong deviations too harshly, as we deemed them to be potentially appropriate for following such things as the degree of engagement of the animal. The augmented loss thus took this form:

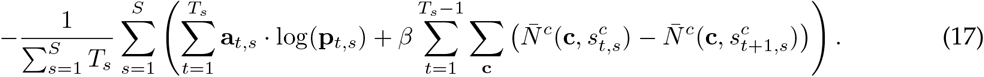

*β* is a hyperparameter to scale this term, which we included in our hyperparameter sweep (covering the values 0.01, 0.1, 1, and 10). Note that the *β* term involves a sum over all possible contrasts **c**, to measure the overall change in the PMF given the current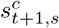, irrespective of the actually presented contrast. The PMFs instantiated by the contrast network are compared in logit space.

The best hyperparameter of the regularised ABCD network was: batch size 16, learning rate initialisation 10^−4^, weight decay 10^−5^, LSTM dimensionality 8, contrast network dimensionality 16.

### Mirror equivariant networks

Some of the learned networks exhibited biases in the two separate arms, which however cancelled out in the ultimate logit sum. This made their outputs hard to interpret. Therefore, we created a network structure which prohibited such asymmetries. We applied this to both the contrast network and the decay network. Neatly, since the one-hot vector of the previous action and the full decay vector have an uneven number of entries, timeout choices and timeout logits automatically get treated separately by such a network. To ensure that the output to, for example, the contrast (0.5, 0), a 50% contrast on the left, was opposite and equal to (0, 0.5), we passed the mirrored input to the network as well:

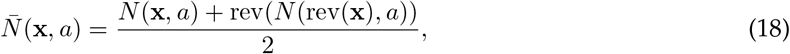

Where we use *N* without a bar to denote a simple MLP, with a single hidden layer, following:

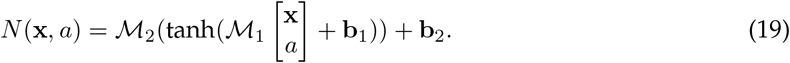

To form the network input, the (optional) scalar *a* gets appended to the vector x. The matrices ℳ and biases *b* are of the appropriate dimensions to map the input first to the hidden units and from there onto the three output logits. We use “rev” to denote the inversion of the order of elements in a vector (e.g. rev((0.5, 0)) = (0, 0.5)). Note that any additional input (the contrast scalar for the contrast network

or previous reward for the decay network) is not part of the inversion. With this formula, inverting the input leads to an inversion of the output logits:

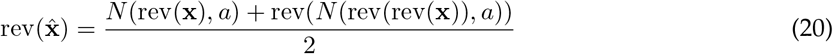

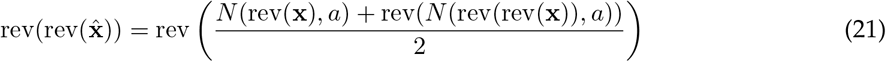

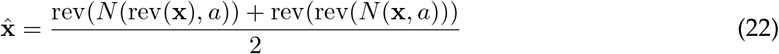

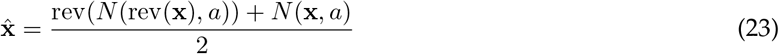

Where we made use of the facts that rev is its own inverse, distributes over addition, and commutes with scalar multiplication. Notably, this also forces the output of the contrast network to be unbiased between leftwards and rightwards choices on a 0% contrast (though the timeout logit is unconstrained).

The division by two is not strictly necessary, as the network could adapt its own weights accordingly, but serves to bring the weights on a comparable scale across architectures. The resulting modulation that was implemented by the contrast scalar was particularly interpretable, as it simply tuned the steepness of the PMF.

## Data availability

We used public data from the IBL. The linked repository contains a script for downloading the exact data.

## Acknowledgments

We thank the Theory working group of the International Brain Lab for discussions and in particular Charline Tessereau for helpful feedback on the manuscript, as well as Jonathan Pillow and Anne Churchland. This work was funded by the Simons Foundation (SAB & PD under ID: 552343, and the IBL in general), the Max Planck Society (SAB, PD), the Wellcome trust (SAB, IBL, PD, ID: 216324), the Alexander von Humboldt Foundation (PD), and Google DeepMind (MKE).

## Authors’ contributions

SAB, MKE, and PD initiated the project. MKE devised the hybrid network framework, which was adapted to this data by SAB, MKE, and PD, and implemented by SAB. The data was collected by members of the IBL (none of the authors were involved). SAB, MKE, and PD performed the data analysis. The paper was jointly written by SAB, MKE, and PD.

## Data and code availability

Data is made available by the IBL, see the documentation at https://docs.internationalbrainlab.org/notebooks_external/data_download.html. Code will be made available upon publication.

## Ethics statement

No ethical approval was required for this study.

## Appendix A Extended Data

## Appendix B Supplemental figures

**Fig. B7.**
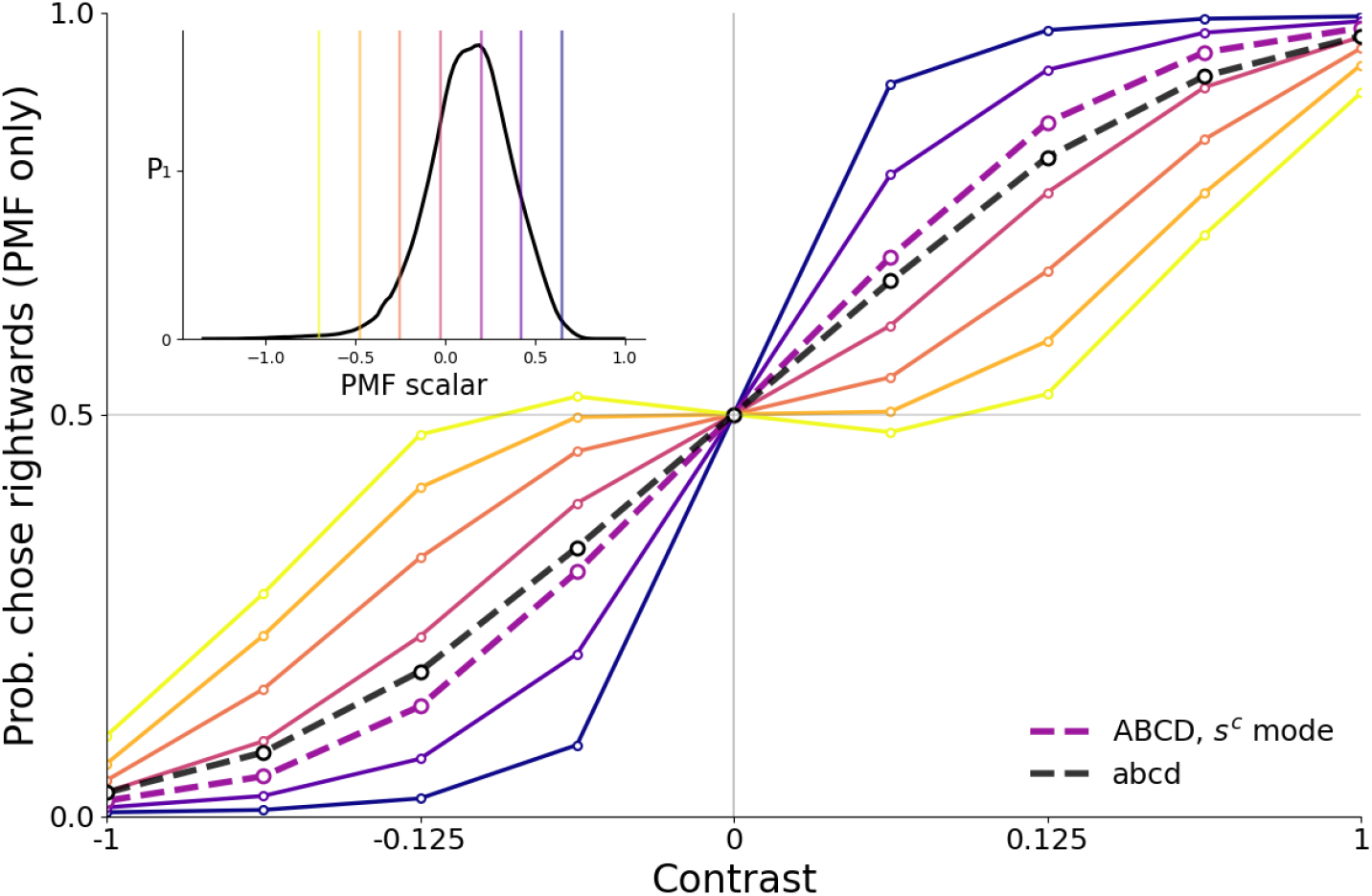
Psychometric functions of ABCD in choice probability space, conditioned on the contrast scalar *s*^*c*^. This is generated by softmaxing the contrast logits by themselves, ignoring history logits (and setting timeout logits to a large negative value), for a visualisation in a more intuitive space. Note that the PMF at the mode of the *s*^*c*^ distribution is more sharply tuned than the static PMF of model abcd.

**Fig. B8.**
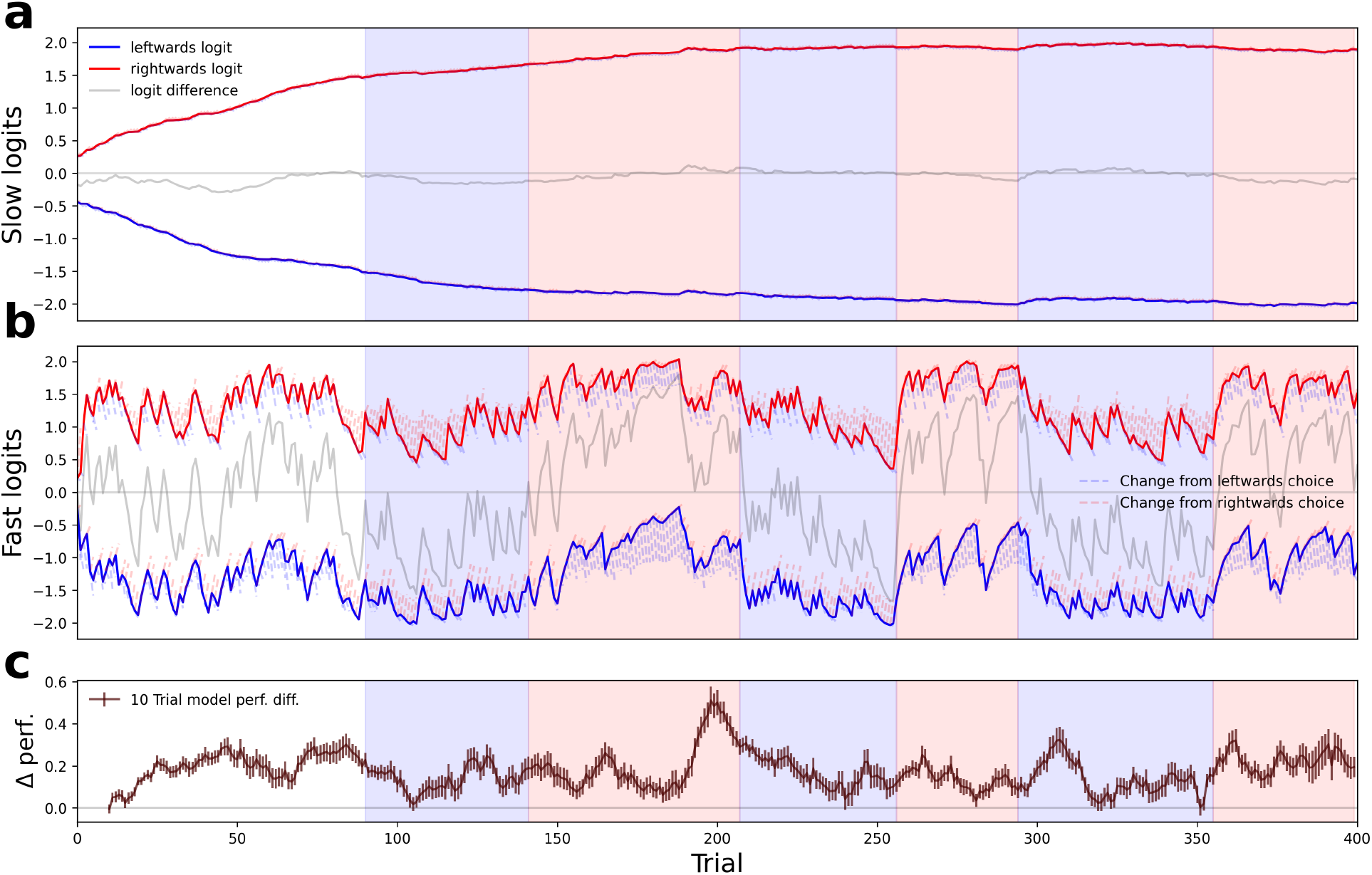
Same logic as 5d-f for sessions of type 2, N=77.

**Fig. B9.**
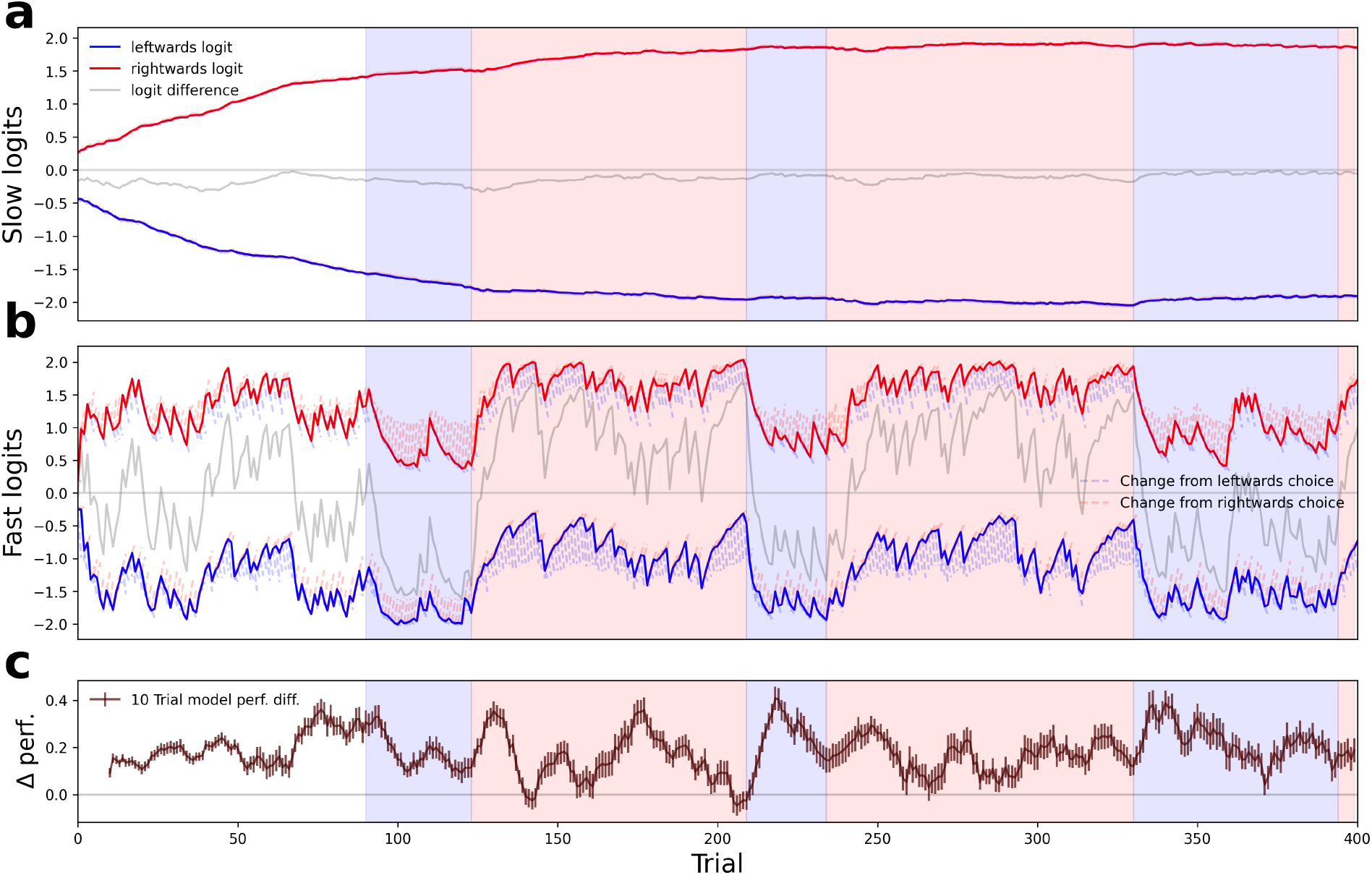
Same logic as 5d-f for sessions of type 3, N=71.

**Fig. B10.**
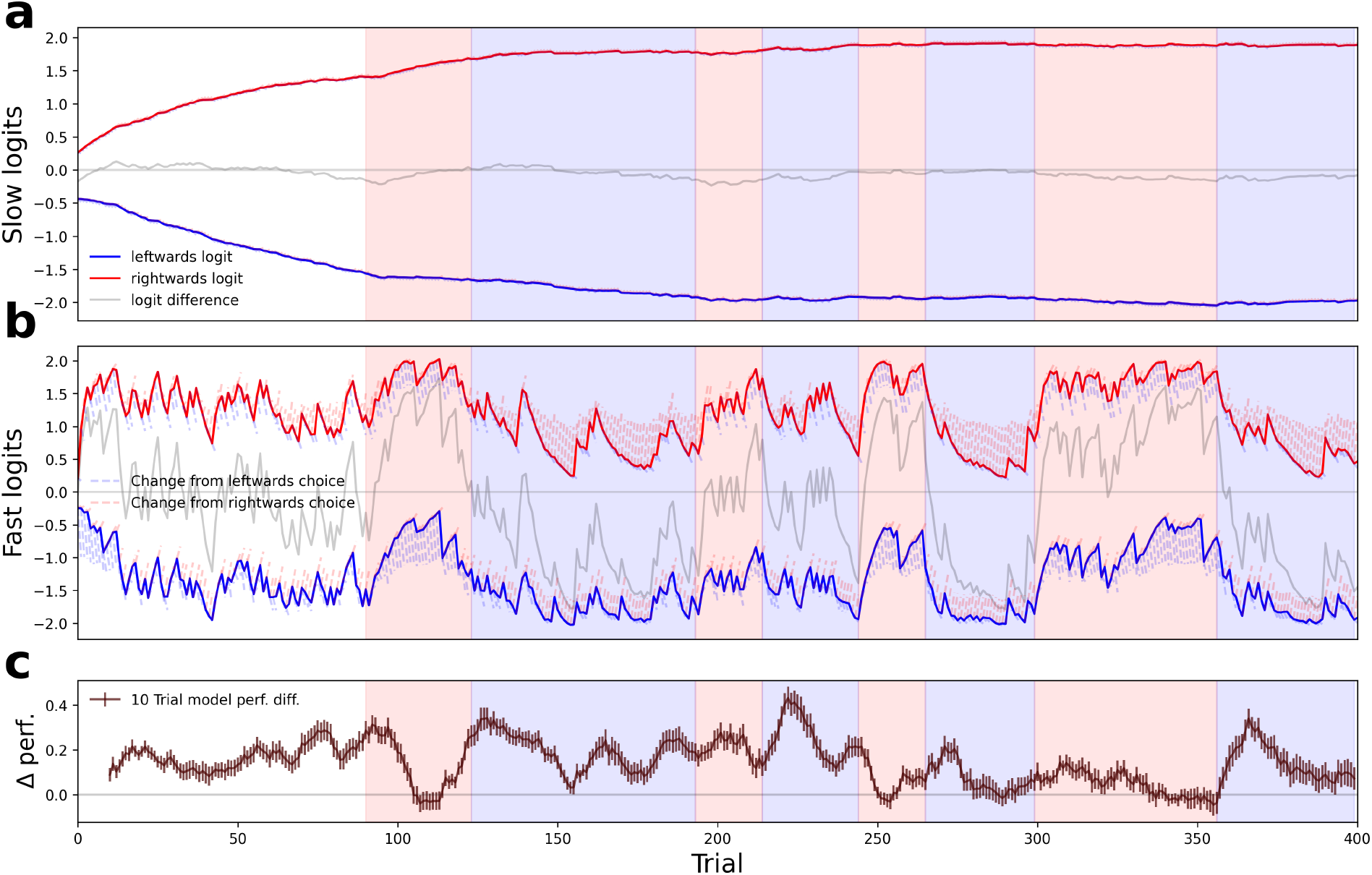
Same logic as 5d-f for sessions of type 4, N=73.

**Fig. B11.**
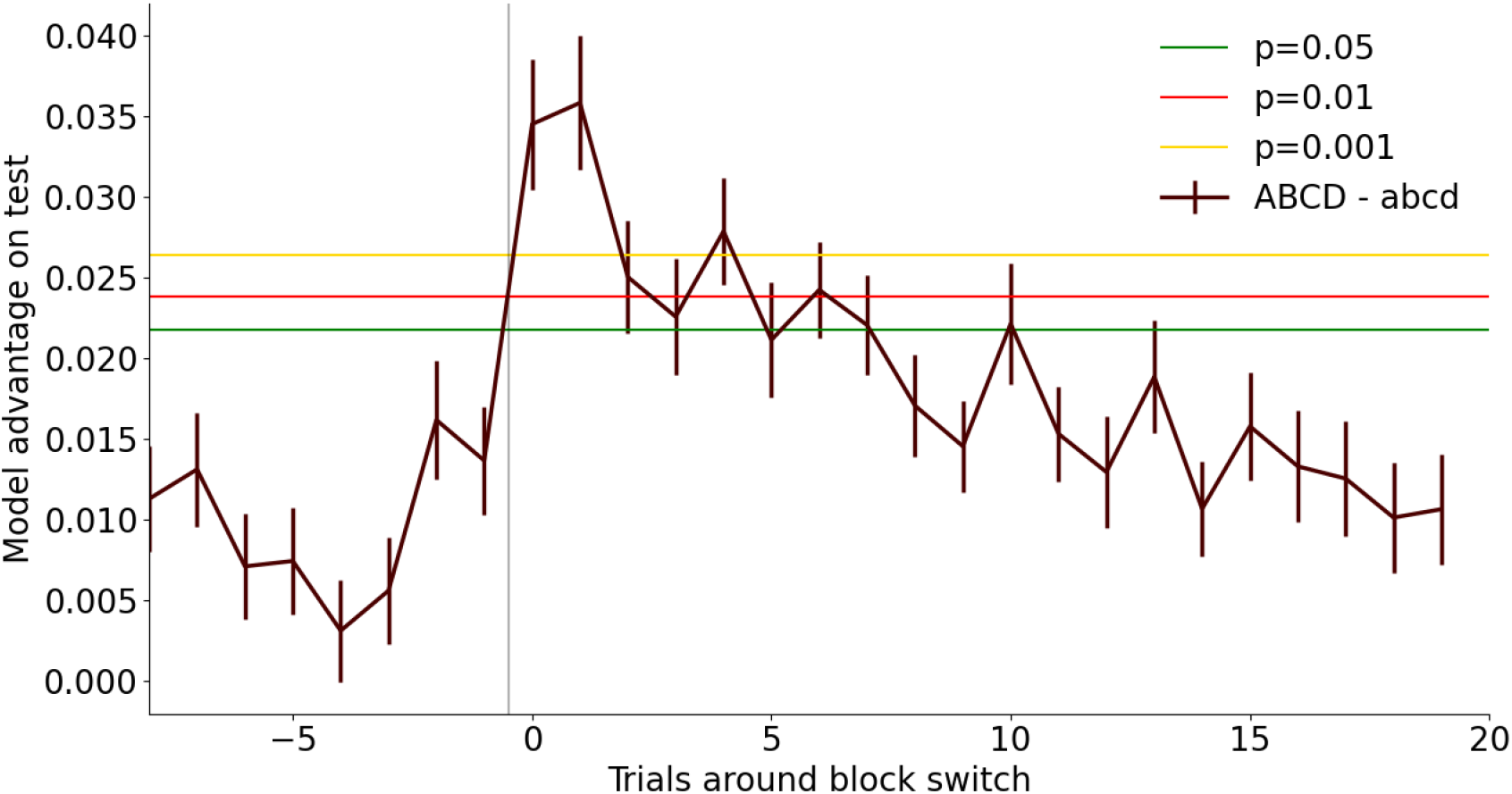
Permutation test of model advantage (difference between the probability assigned to the chosen action by ABCD - assigned probability by abcd) around block switches in the test set. There are 691 block switches in the test set, i.e. every point on the advantage curve is a mean over 691 trials. To construct a null distribution we sample 691 random trials (with replacement) from all available trials (in test sessions), 100,000 times. We can the determine significance at different *α* levels by taking specific quantiles of this distribution (we employ two-sided tests). The model advantage right after a switch is highly significant, before declining somewhat over the next few trials, though staying significant at 0.05 for ∼ 7 trials.

**Fig. B12.**
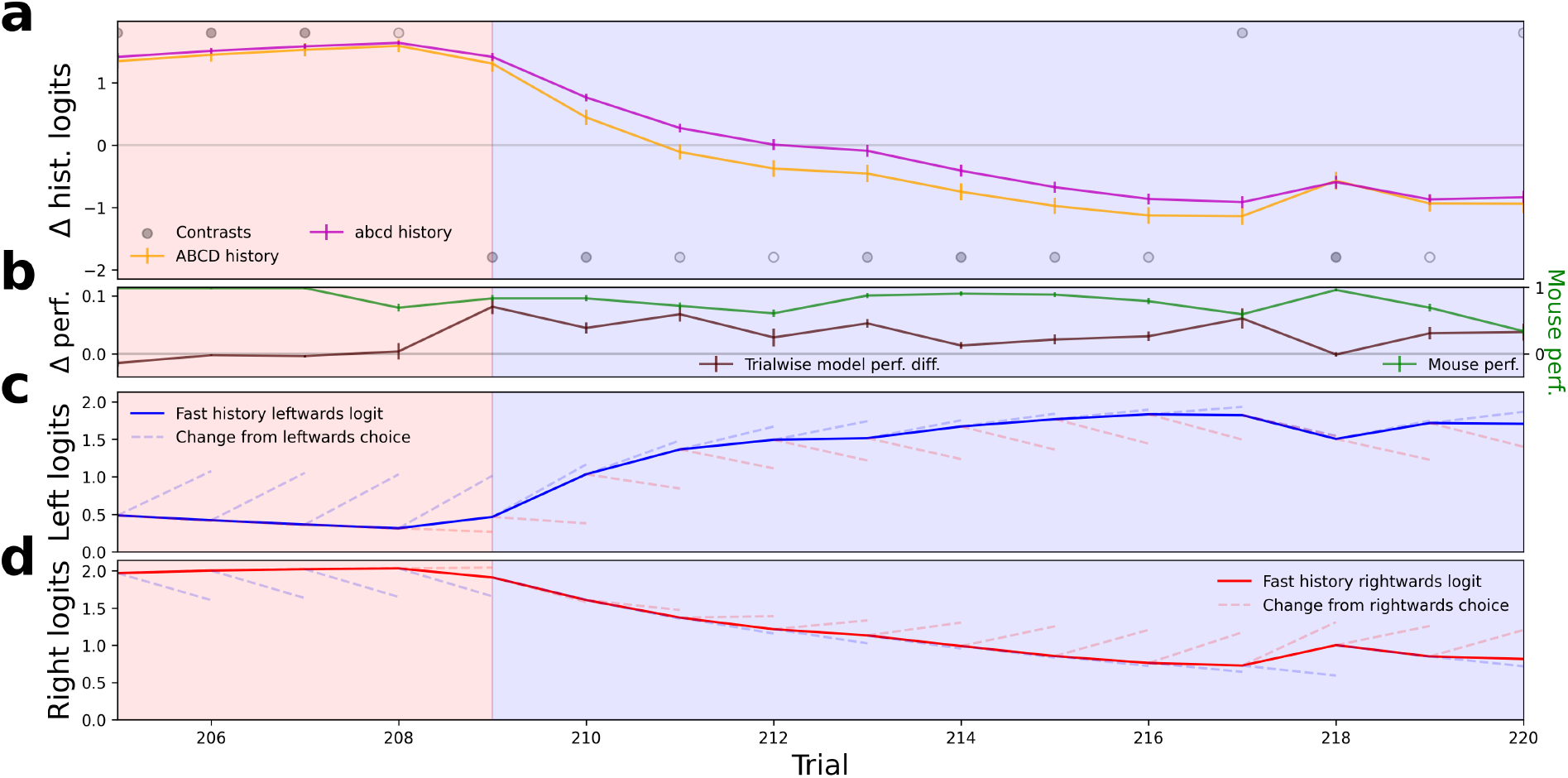
The history term of ABCD reveals that mice can change their bias faster than previously appreciated. This can be seen by zooming in on a small number of trials, showing the workings of ABCD on trials which exhibit the greatest advantage of ABCD over abcd across all sessions of type 3 (N=71). **(a)** Comparison of the mean history term across all animals in this session (the sum of the fast and slow filter in the case of ABCD, errorbars depict ± 1 SEM). Though the models start from a similar point in terms of overall bias, ABCD decreases its bias faster in response to the block change (background colour). **(b)** Trialwise model performance difference (probability assigned to the chosen action by ABCD minus that of abcd) and mouse performance. **(c)** Detailed look at the averaged leftwards logits of the fast history. The whiskers show how the history would change, given a specific choice (the population of mice falls somehwere in between). **(d)** Detailed look at the rightwards logits of the fast history. Note how this history component decreases almost equally in response to correct leftwards choices and incorrect rightwards choices (the whiskers are close together). The stronger decay on unrewarded choices leads to a decrease, even if the choice itself reinforces the bias. Of course, as seen in panel (c), the leftwards logits react differently to the two types of choices.

**Fig. B13.**
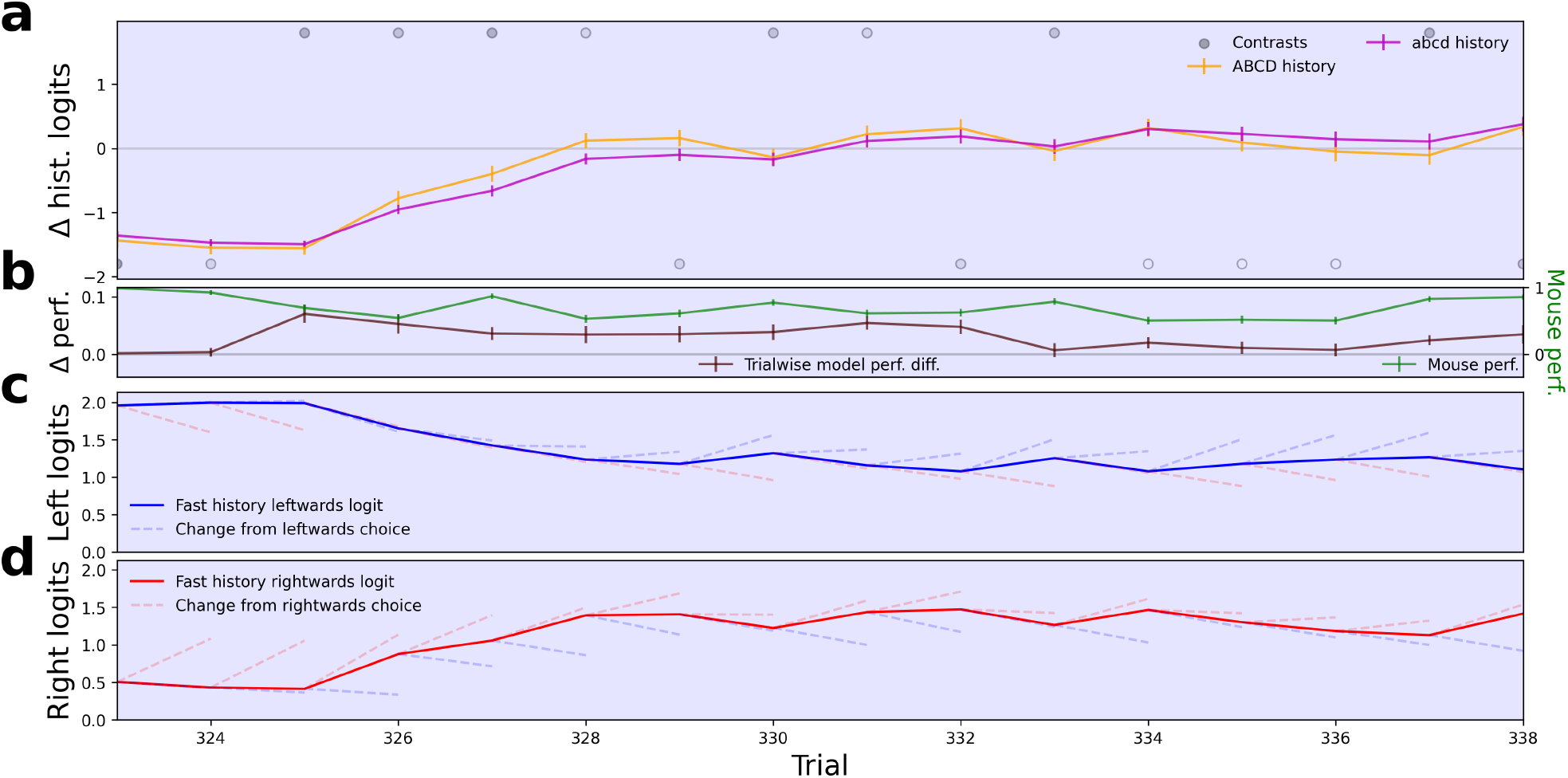
Same logic as B12 for sessions of type 1 (N=74).

**Fig. B14.**
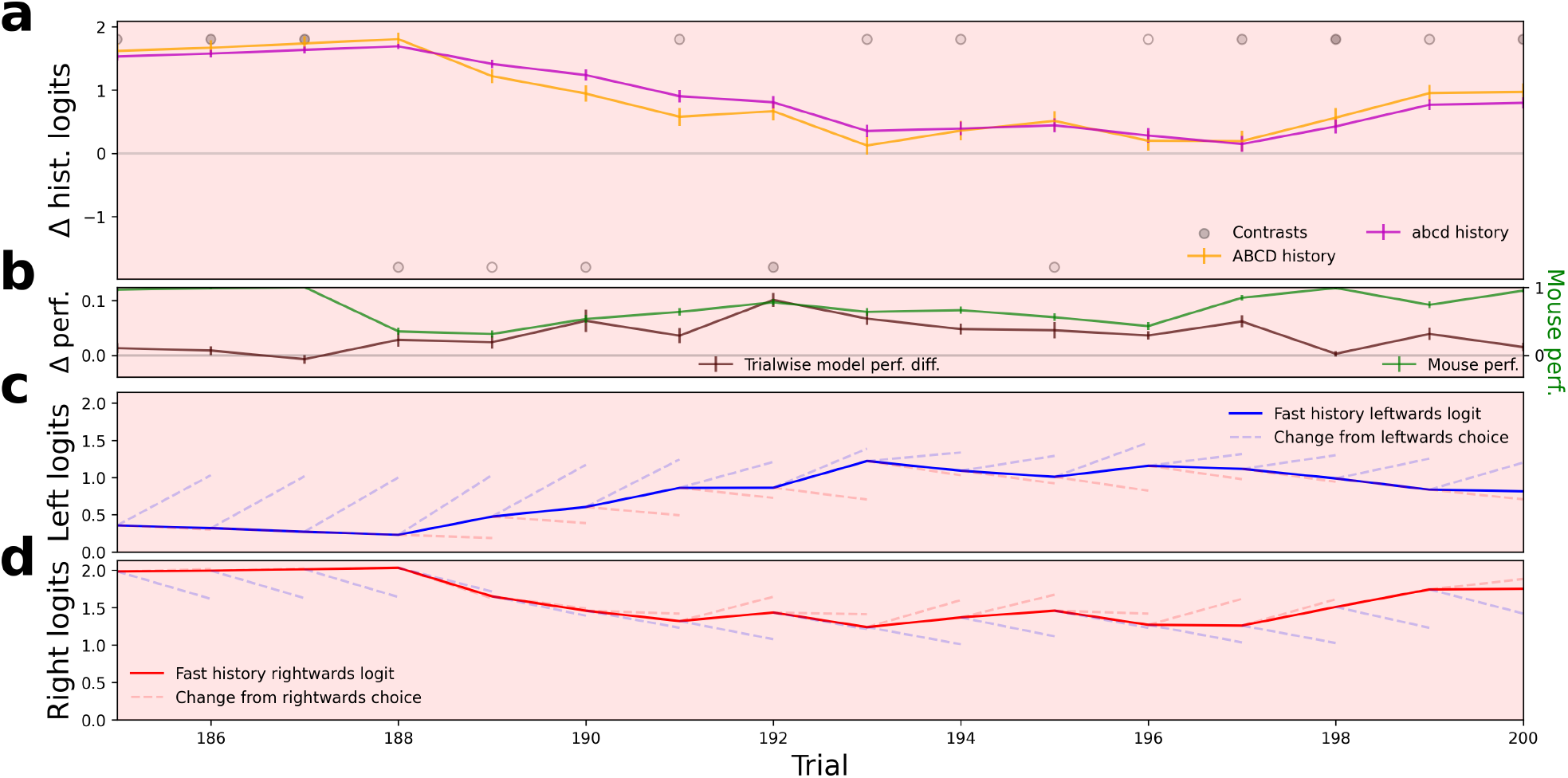
Same logic as B12 for sessions of type 2 (N=77).

**Fig. B15.**
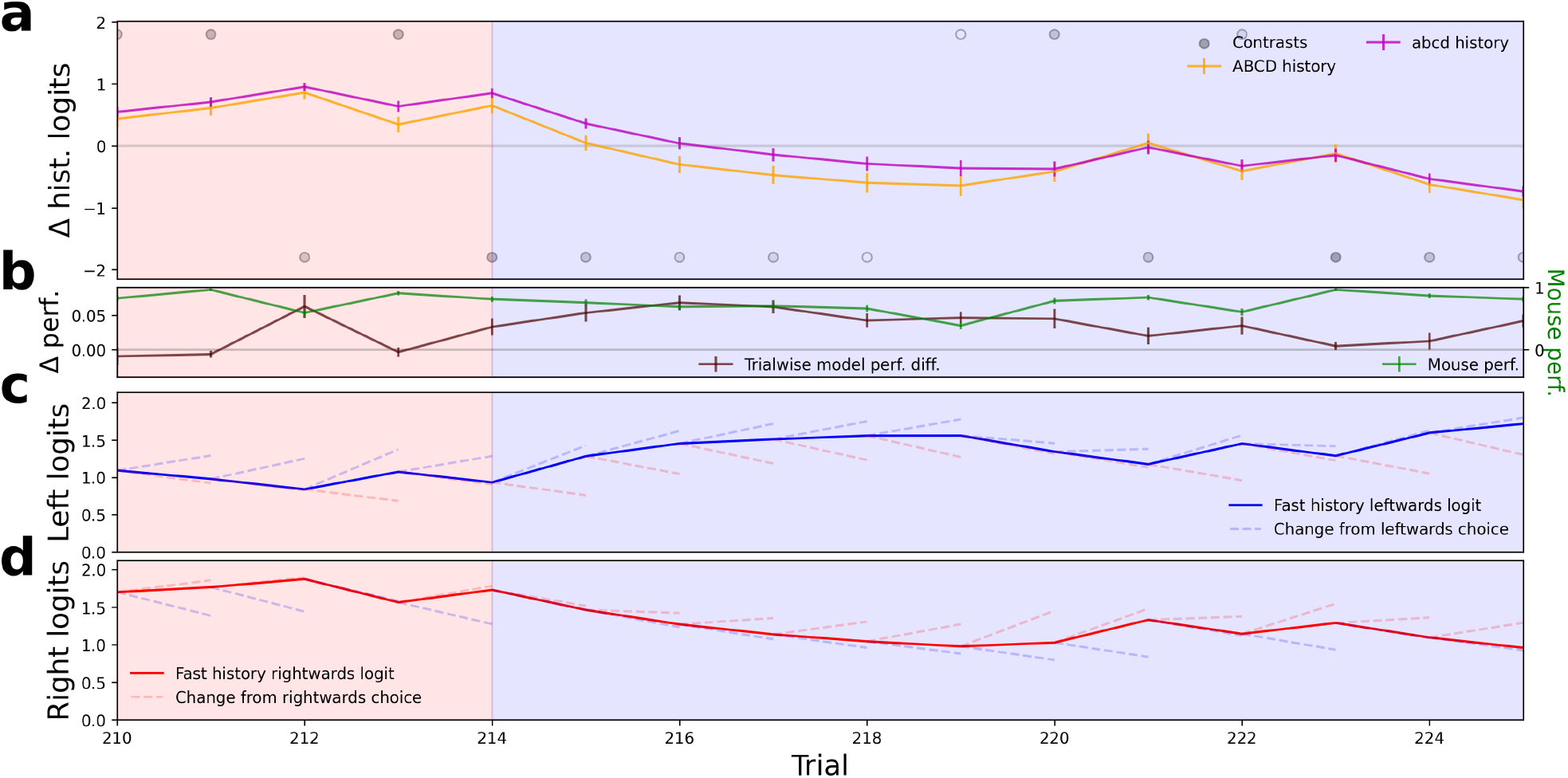
Same logic as B12 for sessions of type 4 (N=73).

## Footnotes

1 Note that there were just 12 distinct “session types” (i.e. predefined sequences of block lengths and stimulus contrasts shown) which mice experienced in order without repeats. This caused the first few session types to be over-represented and introduces patterns into performance trajectories.

2 We use four letters in our model naming scheme, in anticipation of the four extensions which we will apply, using lower case letters to mark the absence of a specific extension and capital letters for their presence.

## References

Ashwood, Z.C., Roy, N.A., Stone, I.R., The International Brain Laboratory, Urai, A.E., Churchland, A.K., … Pillow, J.W. (2022, February). Mice alternate between discrete strategies during perceptual decision-making. Nature Neuroscience, 25(2), 201–212, 10.1038/s41593-021-01007-z Retrieved 2023-05-17, from https://www.nature.com/articles/s41593-021-01007-z

Binz, M., Dasgupta, I., Jagadish, A.K., Botvinick, M., Wang, J.X., Schulz, E. (2024). Meta-learned models of cognition. Behavioral and Brain Sciences, 47, e147,

Bruijns, S.A., Laboratory, I.B., Bougrova, K., Laranjeira, I.C., Lau, P.Y.P., Meijer, G.T., … Dayan, P. (2023). Dissecting the complexities of learning with infinite hidden markov models. bioRxiv, 10.1101/2023.12.22.573001 Retrieved from https://www.biorxiv.org/content/early/2023/12/23/2023.12.22.573001 https://www.biorxiv.org/content/early/2023/12/23/2023.12.22.573001.full.pdf

Castro, P.S., Tomasev, N., Anand, A., Sharma, N., Mohanta, R., Dev, A., … others (2025). Discovering symbolic cognitive models from human and animal behavior. bioRxiv, 2025–02,

Cavanagh, S.E., Hunt, L.T., Kennerley, S.W. (2020). A diversity of intrinsic timescales underlie neural computations. Frontiers in Neural Circuits, 14, 615626,

Dezfouli, A., Ashtiani, H., Ghattas, O., Nock, R., Dayan, P., Ong, C.S. (2019). Disentangled behavioural representations. Advances in neural information processing systems, 32,

Dezfouli, A., Griffiths, K., Ramos, F., Dayan, P., Balleine, B.W. (2019). Models that learn how humans learn: The case of decision-making and its disorders. PLoS computational biology, 15(6), e1006903,

Eckstein, M., Summerfield, C., Daw, N., Miller, K.J. (2024, Jul). Hybrid neural-cognitive models reveal how memory shapes human reward learning. PsyArXiv. Retrieved from osf.io/preprints/psyarxiv/u9ks4

Findling, C., Hubert, F., International Brain Laboratory, Acerbi, L., Benson, B., Benson, J., … Pouget, A. (2023). Brain-wide representations of prior information in mouse decision-making. bioRxiv, 10.1101/2023.07.04.547684 Retrieved from https://www.biorxiv.org/content/early/2023/07/05/2023.07.04.547684.1 https://www.biorxiv.org/content/early/2023/07/05/2023.07.04.547684.1.full.pdf

Funamizu, A., Ito, M., Doya, K., Kanzaki, R., Takahashi, H. (2012). Uncertainty in action-value estimation affects both action choice and learning rate of the choice behaviors of rats. European Journal of Neuroscience, 35(7), 1180–1189,

Gershman, S.J. (2020, November). Origin of perseveration in the trade-off between reward and complexity. Cognition, 204, 104394, 10.1016/j.cognition.2020.104394 Retrieved 2023-06-26, from https://linkinghub.elsevier.com/retrieve/pii/S0010027720302134

Green, D.M., Swets, J.A., et al. (1966). Signal detection theory and psychophysics (Vol. 1). Wiley New York.

Hochreiter, S., & Schmidhuber, J. (1997, 11). Long Short-Term Memory. Neural Computation, 9(8), 1735–1780, 10.1162/neco.1997.9.8.1735 Retrieved from https://doi.org/10.1162/neco.1997.9.8.1735 https://direct.mit.edu/neco/article-pdf/9/8/1735/813796/neco.1997.9.8.1735.pdf

Ji-An, L., Benna, M.K., Mattar, M.G. (2023). Automatic discovery of cognitive strategies with tiny recurrent neural networks. bioRxiv, 10.1101/2023.04.12.536629 Retrieved from https://www.biorxiv.org/content/early/2023/05/03/2023.04.12.536629 https://www.biorxiv.org/content/early/2023/05/03/2023.04.12.536629.full.pdf

Kingma, D.P., & Ba, J. (2017). Adam: A method for stochastic optimization. Retrieved from https://arxiv.org/abs/1412.6980

McCormick, D.A., Nestvogel, D.B., He, B.J. (2020). Neuromodulation of brain state and behavior. Annual review of neuroscience, 43(1), 391–415,

Miller, K.J., Eckstein, M., Botvinick, M.M., Kurth-Nelson, Z. (2024). Cognitive model discovery via disentangled rnns. Proceedings of the 37th international conference on neural information processing systems. Red Hook, NY, USA: Curran Associates Inc.

Nassar, M.R., & Frank, M.J. (2016, October). Taming the beast: extracting generalizable knowledge from computational models of cognition. Curr Opin Behav Sci, 11, 49–54,

Palminteri, S., Wyart, V., Koechlin, E. (2017). The importance of falsification in computational cognitive modeling. Trends in Cognitive Sciences, 21(6), 425–433, https://doi.org/10.1016/j.tics.2017.03.011 Retrieved from https://www.sciencedirect.com/science/article/pii/S1364661317300542

Power, A., Burda, Y., Edwards, H., Babuschkin, I., Misra, V. (2022). Grokking: Generalization beyond overfitting on small algorithmic datasets. Retrieved from https://arxiv.org/abs/2201.02177

Rabiner, L., & Juang, B. (1986). An introduction to hidden markov models. ieee assp magazine, 3(1), 4–16,

Roy, N.A., Bak, J.H., Akrami, A., Brody, C.D., Pillow, J.W. (2021, February). Extracting the dynamics of behavior in sensory decision-making experiments. Neuron, 109(4), 597–610.e6, 10.1016/j.neuron.2020.12.004 Retrieved 2023-05-17, from https://linkinghub.elsevier.com/retrieve/pii/S0896627320309636

Rumelhart, D.E., Hinton, G.E., Williams, R.J. (1986). Learning representations by back-propagating errors. nature, 323(6088), 533–536,

Schaeffer, R., Khona, M., Meshulam, L., Laboratory, I.B., Fiete, I.R. (2020). Reverse-engineering recurrent neural network solutions to a hierarchical inference task for mice. bioRxiv, 10.1101/2020.06.09.142745 Retrieved from https://www.biorxiv.org/content/early/2020/06/12/2020.06.09.142745 https://www.biorxiv.org/content/early/2020/06/12/2020.06.09.142745.full.pdf

Smith, M.A., Ghazizadeh, A., Shadmehr, R. (2006). Interacting adaptive processes with different timescales underlie short-term motor learning. PLoS biology, 4(6), e179,

The International Brain Laboratory, Aguillon-Rodriguez, V., Angelaki, D., Bayer, H., Bonacchi, N., Carandini, M., … Zador, A.M. (2021, May). Standardized and reproducible measurement of decision-making in mice. eLife, 10, e63711, 10.7554/eLife.63711 Retrieved 2023-06-01, from https://elifesciences.org/articles/63711

The International Brain Laboratory, Benson, B., Benson, J., Birman, D., Bonacchi, N., Carandini, M., … Witten, I.B. (2023). A brain-wide map of neural activity during complex behaviour. bioRxiv, 10.1101/2023.07.04.547681 Retrieved from https://www.biorxiv.org/content/early/2023/07/04/2023.07.04.547681.1 https://www.biorxiv.org/content/early/2023/07/04/2023.07.04.547681.1.full.pdf

Thorndike, E.L. (1911). Animal intelligence: Experimental studies. Lewiston, NY, US: Macmillan Press. Retrieved from 10.5962/bhl.title.55072

Wichmann, F.A., & Hill, N.J. (2001, Nov 01). The psychometric function: I. fitting, sampling, and goodness of fit. Perception & Psychophysics, 63(8), 1293–1313, 10.3758/BF03194544 Retrieved from https://doi.org/10.3758/BF03194544

Wilson, R.C., & Collins, A.G. (2019, 11). Ten simple rules for the computational modeling of behavioral data. eLife, 8, e49547, 10.7554/eLife.49547 Retrieved from https://doi.org/10.7554/eLife.49547

